# Latent infection of an active giant endogenous virus in a unicellular green alga

**DOI:** 10.1101/2024.09.03.611062

**Authors:** Maria P. Erazo-Garcia, Uri Sheyn, Zachary K. Barth, Rory J. Craig, Petronella Wessman, Abdeali M. Jivaji, W. Keith Ray, Maria Svensson-Coelho, Charlie K. Cornwallis, Karin Rengefors, Corina P. D. Brussaard, Mohammad Moniruzzaman, Frank O. Aylward

## Abstract

Latency is a common strategy in a wide range of viral lineages, but its prevalence in giant viruses remains unknown. Here we describe the activity and viral production from a 617 kbp integrated giant viral element in the model green alga *Chlamydomonas reinhardtii*. We resolve the integrated viral region using long-read sequencing and show that viral particles are produced and released in otherwise healthy cultures. A diverse array of viral-encoded selfish genetic elements are expressed during GEVE reactivation and produce proteins that are packaged in virions. In addition, we show that field isolates of *Chlamydomonas* sp. harbor latent giant viruses related to the *C. reinhardtii* GEVE that exhibit similar infection dynamics, demonstrating that giant virus latency is prevalent in natural host communities. Our work reports the largest temperate virus documented to date and the first active GEVE identified in a unicellular eukaryote, substantially expanding the known limits of viral latency.

## Introduction

Endogenous Viral Elements (EVEs) are prevalent features in eukaryotic genomes that play key roles in regulation, antiviral defense, and other cellular processes [1–3]. Once linked primarily to integrated retroviruses, it is now recognized that EVEs are derived from a wide range of viral lineages, including ssDNA and dsDNA viruses [4–7]. To date, the largest EVEs discovered are derived from large DNA viruses in the phylum *Nucleocytoviricota,* often called “giant viruses” due to their large genomes and virions. Large EVEs derived from nucleocytoviruses, called Giant Endogenous Viral Elements (GEVEs), are ubiquitous in green algae, brown algae, various fungi, a wide range of other protists, and even some plants and animals [8–11]. GEVEs are prominent features that can contribute large quantities of viral genes to the genomes of their hosts; for example, the genome of the green alga *Tetrabaena socialis* includes two GEVEs totalling >3 Mbp, while the fungus *Rhizophagus irregularis* has the longest contiguously-resolved GEVE at 1.5 Mbp [8,9].

Despite the large contribution of GEVEs to many eukaryotic genomes, it remains unknown whether these elements are derived from active integration of nucleocytoviruses as part of their infection cycle or merely accidental integration that occurs during stalled infections. Interestingly, studies dating back as far as the 1970s have observed the formation of icosahedral particles from otherwise healthy cultures of protists, but it has remained unclear if this can be attributed to the reactivation of latent viruses or other factors such as persistent infection [12,13]. The best-studied example of a putatively active GEVE to date is in the multicellular brown alga *Ectocarpus* sp. *7*, where a 330 kbp endogenous nucleocytovirus has been linked to virus-like particle (VLP) formation in reproductive tissues [14–16], but even here the specific activity of the GEVE has not been directly shown. Indeed, the viability of many GEVEs is questionable, and many appear to be silenced through methylation and chromatin remodeling, while others have undergone large-scale erosion and genomic rearrangements that likely led to their inactivation [9,10,17].

Given the recent widespread discovery of GEVEs in protist genomes [8,10,18], it is important to determine if these elements arise from a viral infection strategy involving latency and genome integration. To shed light on the activity of GEVEs and their potential for virion production, we studied the model green alga *Chlamydomonas reinhardtii*, which has been used for decades in detailed analyses of cilia, photosynthesis, and other aspects of eukaryotic biology [19,20]. The recent observation that some field isolates of *C. reinhardtii* harbor signatures of GEVEs suggests that this alga may also be a useful system for in-depth analysis of endogenous giant viruses [21]. We used a combination of long-read sequencing, transcriptomics, proteomics, and additional surveys of field isolates to examine the activity of GEVEs in *C. reinhardtii* and their role as part of the latent infection cycle of giant viruses. Our work describes the largest known latent virus and the first known virus to infect the model green alga *C. reinhardtii,* thereby highlighting the importance of latency in host-virus interactions in the environment.

## Results and Discussion

### Long-read sequencing resolves a contiguous GEVE

Firstly, we used long-read Oxford Nanopore sequencing to obtain a high-quality draft assembly of *C. reinhardtii* strain CC-2937 (see Methods for details). This strain was selected because a previous study using short-read sequencing found that it contained the most GEVE signatures among all *C. reinhardtii* strains surveyed [21]. We recovered a high-quality assembly with an estimated genome size consistent with the latest *C. reinhardtii* reference genome [22]. We screened the polished contigs of the assembly for nucleocytovirus signatures using ViralRecall [23] and recovered a 617 Kbp GEVE flanked by eukaryotic sequences within a 2.8 Mbp contig (Fig. 1A). Other than the GEVE region, the contig corresponds to chromosome 15 of the latest *C. reinhardtii* CC-4532 assembly. The GEVE was contiguous and delimited by terminal inverted repeats (TIRs) 10.8 and 14.8 kbp in length, with the difference attributable to a variable-length satellite array present in the TIRs. This GEVE is almost twice as long as was previously estimated using short-read sequencing [21], underscoring the importance of long-read sequencing to accurately delineate large endogenous viral elements.

**Fig. 1.**
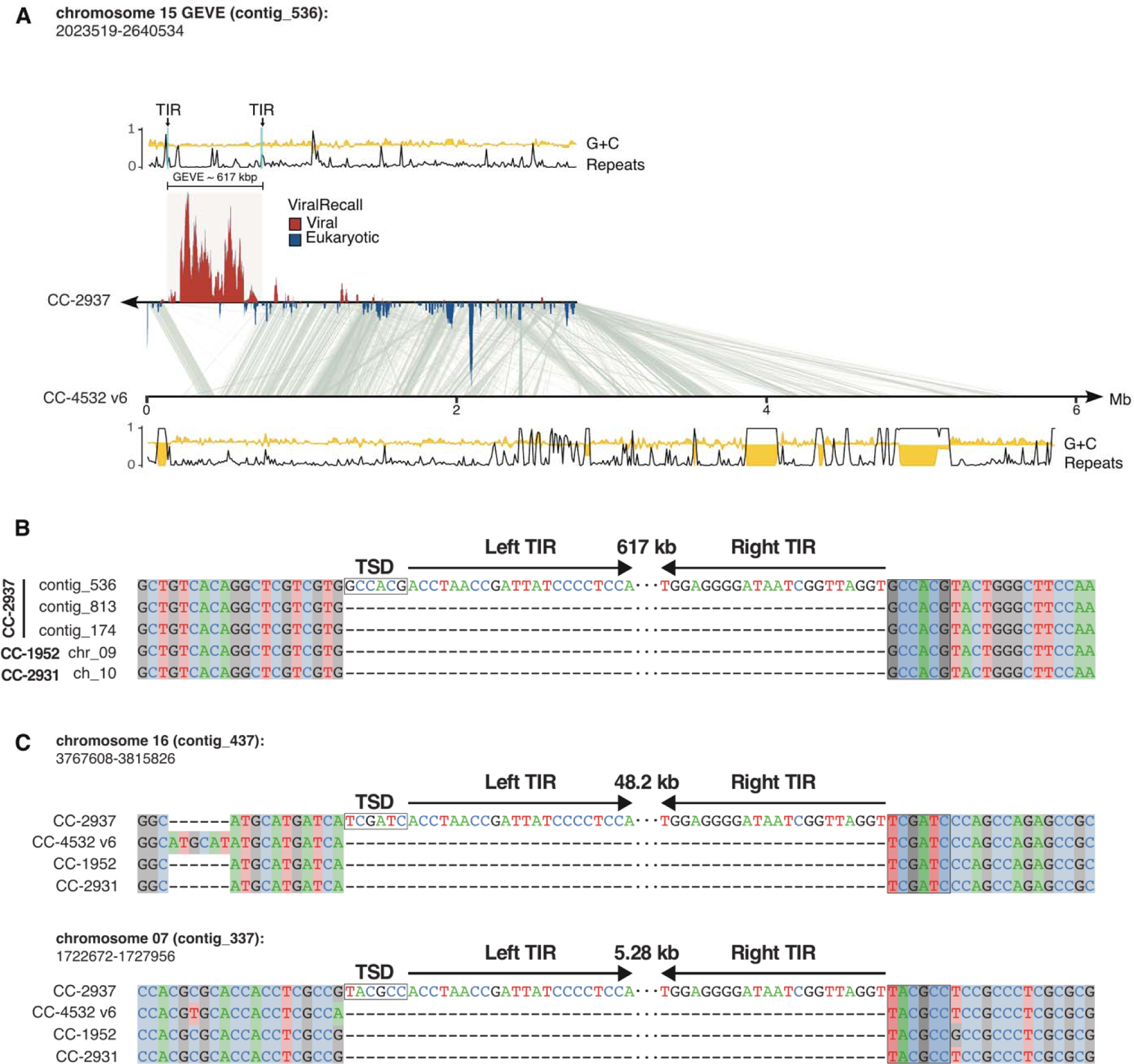
Features of the *C. reinhardtii* CC-2937 GEVE and its insertion site. **(A)** Line plots representing the tandem repeats (black) and GC fraction (yellow) of the viral contig_536. The TIRs flanking the GEVE are marked and shown with arrows. Regions in the contig with viral signatures are represented by ViralRecall scores >0 (red), while eukaryotic regions represent scores <0 (blue). Synteny blocks between the viral contig and chromosome 15 of the reference genome (*C. reinhardtii* CC-4532 v6) are shown in gray. The tandem repeats and GC fraction tracks are also shown for this chromosome at the bottom. **(B)** Alignment of five independent copies of the interspersed repetitive element in which the GEVE is integrated. Only the ends of the TIRs are represented and the 6 bp TSD is highlighted by a box. **(C)** Integration sites of two putative GEVE relics on chromosomes 16 and 7. The TIRs and TSDs of the insertions in CC-2937 are shown relative to three divergent strains that do not carry the insertions.

We predicted 579 ORFs (data S1) from the GEVE that include a complete set of *Nucleocytoviricota* hallmark genes, such as family B DNA polymerase (PolB), two double-jellyroll major capsid proteins (MCPs), multi-subunit RNA polymerase homologs, an A32 packaging ATPase, and a VLTF3 transcription factor [24]. The %GC content of the GEVE was only slightly lower than the flanking regions (60.6% versus 62.8%), and also slightly below the genome-wide %GC content reported for *C. reinhardtii* (64%) [22,25]. Many GEVEs show a clear deviation in nucleotide composition compared to the genomes of their hosts [8], but we found no detectable discrepancy (fig. S1).

To determine the integration site of the GEVE, we compared our CC-2937 assembly with the genomes of the reference strain CC-4532 and two field isolates, CC-1952 and CC-2931 [26]. We mapped the TIRs to an intergenic region downstream of the Cre15.g635700 gene. This exhibits significant structural variation, and the sequence flanking the TIRs corresponds to an ∼9 kbp interspersed repetitive element absent from the other strains at this locus. Independent copies of this repeat are found in CC-2937 at two other regions on chromosomes 4 (contig_813) and 3 (contig_174), and at single loci in CC-1952 (chromosome 9) and CC-2931 (chromosome 10). By aligning these five repeat copies, we determined the exact insertion site and TIR boundaries of the GEVE (Fig. 1B). The TIRs feature the terminal motif “ACC-GGT” and are flanked by a 6 bp target site duplication (TSD).

We identified sequences homologous to the GEVE TIRs at two other regions in the CC-2937 genome, on contig_437 (chromosome 16) and contig_337 (chromosome 7). Comparison to the other strains revealed that these sequences also correspond to insertions unique to CC-2937, although, unlike the GEVE, the flanking sequences are non-repetitive, and the signatures of integration can be directly resolved (Fig. 1C). Insertions in chromosomes 16 and 7 were 48.2 kbp and 5.28 kbp long, respectively. The termini of these insertions perfectly match the left and right ends of the GEVE TIRs, and both are flanked by distinct 6 bp TSDs. Most of the integrated sequences can be mapped to regions of the GEVE, suggesting that they represent relics of closely related viruses that have undergone deletion following endogenization. We also found relics of TIR sequence, several of which were flanked by 6 bp TSDs, among the other available *C. reinhardtii* genomes (table S1). Altogether, the widespread presence of GEVE relics in other strains as well as other locations in the CC-2937 genome suggests that viral integration is a common occurrence, and that there is likely a strong selection for large mutations to deactivate GEVEs.

TSDs of fixed lengths are associated with distinct families of DD(E/D) integrase enzymes, which introduce staggered nicks in the target DNA that, following repair, result in duplications that correspond to the length of the stagger between the two DNA strands [27]. The GEVE carries several integrase genes, although they all belong to the *IS630-Tc1-Mariner* superfamily that introduces “TA” dinucleotide TSDs, and they are encoded by virus-specific selfish elements (see below). The 6 bp TSDs are associated with specific members of a broad assemblage that includes retroviral-like integrases (including integrases from LTR retrotransposons and Polintons) and integrases of mobile elements, including *IS3* and *IS481* [28]. We did not detect any candidate integrase that would produce 6 bp TSDs in the GEVE, and there is also no prior evidence of these enzymes being associated with the ‘ACC-GGT’ termini. Nonetheless, the presence of 6 bp TSDs flanking three independent endogenization events implies active integration by specific, but yet uncharacterized, enzymatic machinery. Nimaviruses that infect crustaceans have acquired retroviral-like integrases that they use for integration [29], and it is possible that a divergent integrase was acquired in a similar manner here.

### Viral particles are produced in C. reinhardtii CC-2937 cultures

Next, we sought to assess whether this viral element was active and could produce viral particles in cultures of *C. reinhardtii* CC-2937. We monitored virion production in cultures from inoculation to stationary phase by performing a PCR assay targeting the viral *mcp* gene on 0.45□µm-filtered supernatants treated with DNAse to eliminate non-encapsidated host DNA (see Methods for details). Our results indicated that free virions began to accumulate as the cultures reached the stationary phase at six days post-inoculation (avg. 1.0 × 10^7^ cells mL^-1^) (Fig. 2A).

**Fig. 2.**
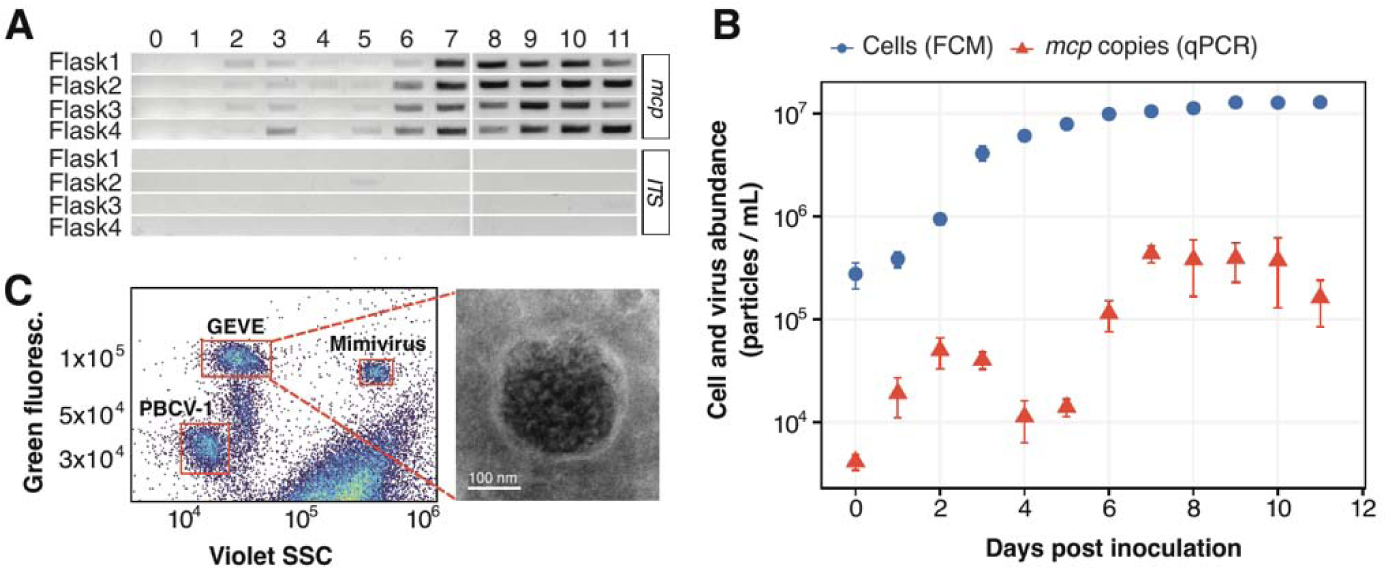
Late exponential cultures of *C. reinhardtii* CC-2937 show evidence of virion production. **(A)** Gel image of a PCR assay targeting the viral major capsid protein (*mcp*) gene and host ITS region. The assay was performed on DNAse-treated supernatants from four culture replicates, sampled daily for 11 days. **(B)** Quantification of viral and host cell abundances over 11 days in four culture replicates. Viral abundance in the supernatants was measured by qPCR (red), while host cell density was assessed using flow cytometry (FCM) (blue). Error bars represent the standard deviation. **(C)** Flow cytometry analysis (left) of concentrated supernatants alongside two positive controls of known large DNA viruses mixed in with the sample. The right panel displays an electron micrograph of the concentrated viral fraction of a 9-day-old culture, showing a virus-like particle (VLP) identified through negative staining.

To examine this trend in more detail, we quantified viral DNA in these samples through qPCR by targeting the GEVE *mcp* gene sequence and comparing the amplification results to a calibration curve generated from amplifying the *mcp* sequence in a DNA construct of known concentrations (fig. S2, details in Methods). The production of virions in culture occurred in two waves, peaking at day two and day seven post-inoculation with an average of 5.0 × 10^4^ and 44 × 10^4^ *mcp* copies mL^-1^, respectively (Fig. 2B). The cultures appeared healthy and did not crash, demonstrating that a virulent infection did not take place. These results indicate that low levels of viral production were maintained at high host cell densities, leading to a ratio of virions to host cells of ∼0.05:1. We also verified the presence of free viral particles using flow cytometry of the viral-fraction material concentrated via tangential flow filtration (details in Methods). We identified a distinct population of particles with comparable staining signature as large dsDNA nucleocytoviruses (Fig. 2C, fig. S3 A-C) (positive controls taken along in our analysis) [30]. Lastly, negative stain electron microscopy of concentrated CC-2937 supernatants consistently showed spherical particles ∼200 nm in diameter with electron-dense cores (Fig. 2C, fig. S4).

The presence of viral particles confirms that the GEVE is active, and it is therefore appropriate to coin a name to refer to this novel viral isolate. For the species taxon we propose the binomial name *Punuivirus latens*. The genus name draws inspiration from the Incan mythology deity Puñuy, who is linked to dreams and the act of sleeping, while the species name refers to the latent infection strategy of this virus. For the viral isolate we use the trivial name Punuivirus cr2937.

### High transcriptional activity of GEVE genes during activation

To examine the viral infection cycle in more detail, we grew duplicate cultures over a 7-day time course and harvested cells at different time points within the growth cycle to assess transcript abundance using deep-sequenced RNA-Seq (fig. S5). Consistent with our qPCR results, we found that the expression of viral genes peaked at the late exponential and early stationary phases of host growth (∼6 days post-inoculation, 4.5 - 5.4 × 10^6^ cells mL^-1^) (Fig. 3A). Almost all GEVE genes were expressed during at least one point along the time-course (n=499, 86%), including the complete set of NCLDV markers (Fig. 3B), demonstrating full activation of viral gene expression (data S2). We used self-organizing maps to demarcate the genes into two distinct clusters, based on whether they were primarily expressed before peak viral production (BVP cluster; days 3-4, n=24 genes) or during peak viral production (DVP cluster; days 5-7, n=167 genes; Fig. 3C). Genes expressed before peak viral production tended to be co-localized near the ends of the GEVE and within the TIRs (in seven clusters of at least two genes, with only two not being colocalized with another gene), while the central region was populated mostly by genes expressed during peak viral production (Fig. 3A). The early expression of the BVP cluster prior to peak viral production, together with the colocalization of many of these genes on the GEVE, suggests that these may have a potential role in the suppression of viral activation. DESeq2 analyses revealed that approximately one-third of the GEVE genes (191 out of 579) were differentially expressed (Fig. 3D, fig. S6). Transcripts enriched during peak viral production include the *mcp* and other structural genes needed for virion formation, consistent with the activation of these genes during virion biogenesis.

**Fig. 3.**
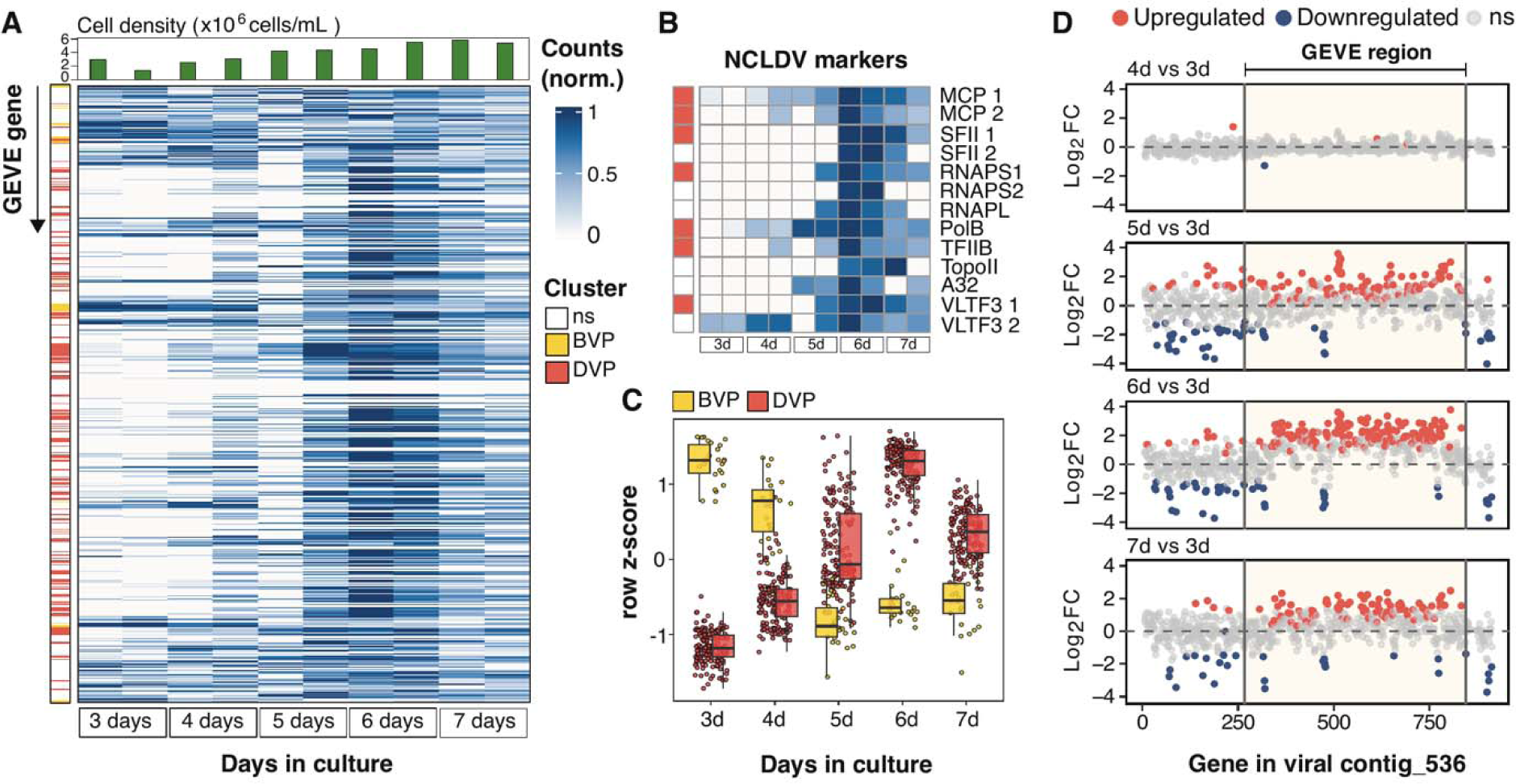
RNA-Seq results of *C. reinhardtii* CC-2937 cultures at different stages of growth. **(A)** Heatmap of min-max normalized counts for each GEVE gene, obtained from culture flasks inoculated 3-7 days prior to collection date. Each row corresponds to a GEVE gene, ordered as they appear along viral contig_536. Each pair of columns represents biological replicates of a specific time point. Row annotations indicate the assigned cluster based on gene expression patterns (see Methods for details): before peak viral production (BVP, yellow), and during peak viral production (DVP, red). Non-significantly differentially expressed genes (ns) were excluded from the clustering analysis. Column annotations show cell counts on the day of sampling (top panel). **(B)** Heatmap of min-max normalized counts specifically for hallmark NCLDV genes [24]. Row annotations indicate the assigned cluster. **(C)** Gene expression patterns of differentially expressed genes (DEGs), colored by their assigned cluster using self-organizing maps. **(D)** Shrunken Log_2_-fold expression changes (Log_2_FC) of genes along the viral contig, as determined by DESeq2. Day three serves as the reference for all comparisons. Each dot represents a gene in the same position as it appears on contig_536. The horizontal dashed line marks no change in gene expression (Log_2_FC = 0).

### Prevalence of GEVE-encoded selfish genetic elements

The GEVE encodes a number of selfish genetic elements, including a single *Metaviridae* LTR retrotransposon, at least seven putative homing endonuclease genes (HEGs), and several Fanzor-encoding elements. All these elements were expressed in our RNA-Seq time-course, demonstrating that these selfish genetic elements are active. The GEVE-encoded HEGs include five inteinic LAGLIDADG nucleases and two freestanding HNH-3 endonucleases. The inteinic LAGLIDADG HEGs are located within the RNA polymerase alpha subunit (RNAPL) (n=2), RNA polymerase beta subunit (RNAPS) (n=2), and the DNA polymerase family B (PolB) (n=1) genes (fig. S7A). The freestanding HNH-3 endonucleases are located proximal to the GEVE’s major capsid gene (fig. S7B). The LTR retrotransposon belongs to the family *Gypsy-4_cRei*, which introduces 5 bp TSDs, and is present at several locations in the *C. reinhardtii* genome (data S3).

The GEVE-encoded Fanzor elements can be split into three distinct families that we refer to as A, B, and C, with copies from each family being nearly identical (>99% nt). For each family, the full-length element also includes a gene encoding a *IS360-Tc1–Mariner* transposase. Within the GEVE, we observed three full length copies of family A, five of family B, and four of family C. In addition to full length elements, we also observed other arrangements including elements that consisted of only the Fanzor gene and right-end guide, non-autonomous transposons that had both ends maintained but gene content was absent or highly degraded, and other element fragments (fig. S8A, data S4). We were able to find homologs to our Fanzor proteins encoded in the GEVEs of other green algae, and we constructed a phylogeny of all these elements together with other references (fig. S8B). Fanzor families A and B are related, while C belongs to a distinct lineage. All of the *C. reinhardtii* GEVE Fanzors belonged to the previously-defined Fanzor 1 lineage that is associated with diverse mobile elements in eukaryotic and giant virus genomes [31]. Moreover, we found a Fanzor fragment within chromosome 17 of the *C. reinhardtii* reference genome (strain CC-4532) that bore 80% nucleotide identity to the sequence from Fanzor C. Together with the apparent mobility of the viral-encoded LTR retrotranposon, these findings demonstrate widespread sharing of selfish genetic elements between virus and host, indicating that endogenous giant viruses are important vectors of selfish DNA in eukaryotes.

### Proteins encoded by selfish genetic elements are packaged into virions

To confirm the presence of free virions, and identify the suite proteins that are likely packaged, we performed liquid chromatography–tandem mass spectrometry (LC-MS/MS) on the supernatants of aging *C. reinhardtii* CC-2937 cultures. A total of 43 proteins were identified with high confidence (at least two Peptide Spectrum Matches in distinct samples; see Methods and data S5). Among these, the MCP was by far the most abundant protein detected, as expected for free virions (Fig. 4). Other abundant proteins included a pectate lyase likely associated with host cell wall degradation [32], a putative envelope protein, both multi-subunit RNA polymerase subunits, several putative viral helicases, DNA topoisomerase II, and a putative procollagen galactosyltransferase, all of which have been found to be packaged in other nucleocytoviruses [32–34]. Most of the other packaged proteins had no predicted function. Based on the GEVE particle diameter (∼200 nm) this number of encoded proteins is within the expected trends observed for other nucleocytoviruses with similar virion size, including *Emiliania huxleyi* Virus 86 (EhV), *Aureococcus anophagefferens* Virus (AaV), *Marseillevirus*, and *Melbournevirus* [33]. Our proteomic analysis detected group B and C Fanzors in the virions, as well as the Gag protein from the LTR retrotransposon (Fig. 4). To our knowledge, this is the first example of the packaging of proteins from mobile elements into viral capsids, and it suggests that the effectors are active immediately upon cellular entry during viral infection. Work in bacteriophages has shown that HEGs can mediate inter-viral competition during co-infection [35,36], suggesting that Fanzors and other selfish genetic elements encoded in GEVEs may also play a similar role.

**Fig. 4.**
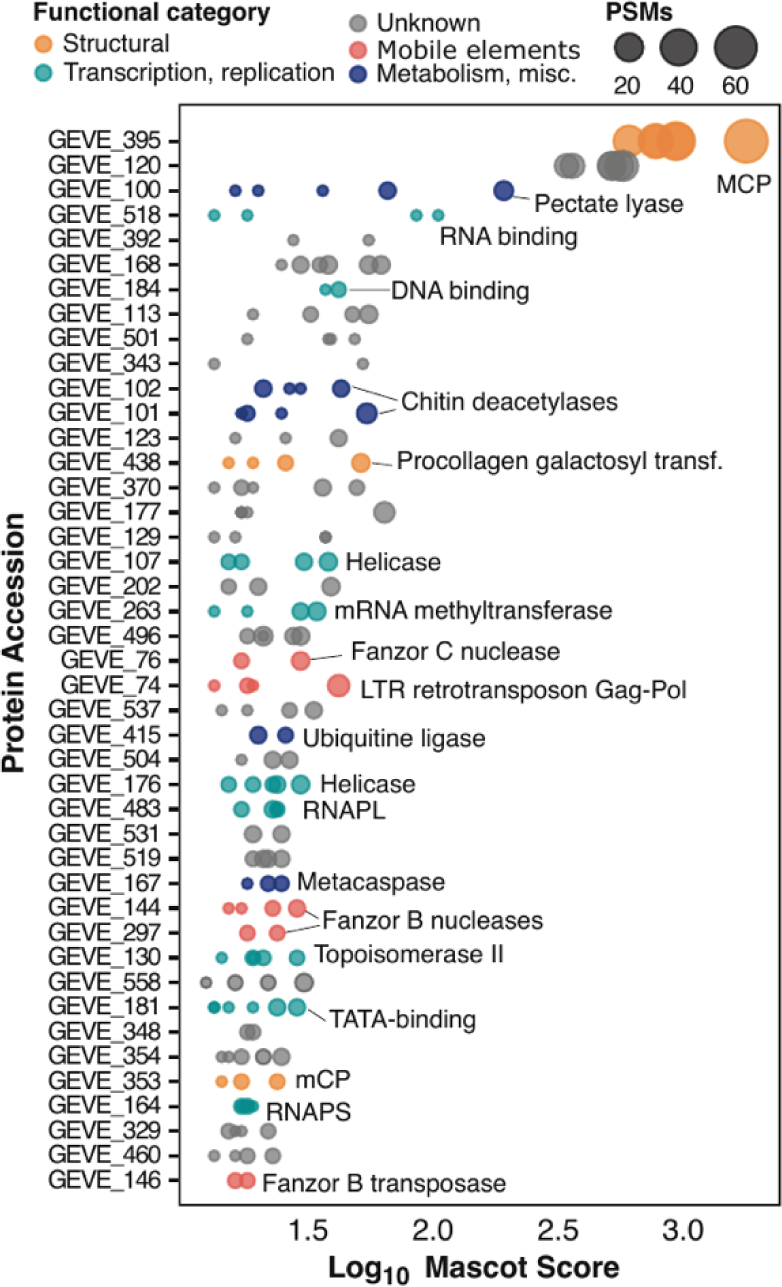
Proteomic analysis of GEVE virions from concentrated supernatants of *C. reinhardtii* CC-2937. Peptides were identified using LC-MS/MS across three biological replicates, each analyzed in duplicate. Each dot represents a protein identified in a technical replicate, with dot size indicating the number of Peptide Spectrum Matches (PSMs) to reflect relative protein abundance, and dot color representing the predicted functional category. Proteins are arranged based on their Mascot database search scores.

### Prevalent latent giant viruses in *Chlamydomonas* populations of Swedish lakes

To assess the prevalence of latent viral infection in a distinct natural population of *Chlamydomonas* sp., we analyzed monoclonal culture strains isolated in 2016 from Örsjön and Krageholmssjön, two lakes situated in southern Sweden. These isolates fall within the *Chlamydomonas* genus and are closely related to *C. reinhardtii* as indicated by molecular analysis of 18S rRNA gene amplicons (fig. S9). Thirteen of the 18 isolates (72%) from Örsjön, and twelve of the 20 from Krageholmssjön (60%) tested positive for amplification of nucleocytovirus *mcp* genes (fig. S10, table S2), indicating a prevalence of latent viruses in this population. These monocultures all have continued to grow well in the laboratory and have not undergone any crashes, similar to CC-2937, demonstrating that the virus is not lethal. To identify the viruses associated with these strains, we sequenced the *mcp* genes from isolates from Örsjön and Krageholmssjön (Ors24 and Kgh18, respectively). Phylogenetic analysis confirmed that these isolates are related to GEVEs previously found in green algal genomes (fig. S11). In addition, we performed low-coverage PacBio sequencing on strain Ors24 that yielded a viral DNA polymerase B sequence, and phylogenetic analysis of this gene confirmed the placement of this isolate within a GEVE clade (fig. S12).

We selected strain Ors24 for thin-section TEM at different growth stages to investigate if viral particles could be observed. Consistent with our findings of *C. reinhardtii* strain CC-2937, we observed viral particles ∼225 nm in diameter that appeared primarily in the mid-exponential phase (Fig. 5A). We observed virions in up to 3% of the cells (fig. S13); because virions would only be expected to be visible in the later stages of a lytic infection program, this suggests that a larger fraction of cells was undergoing active viral infection at that time. Virions with clear icosahedral symmetry were formed from apparent virus factories (Fig. 5B-C), indicating that these structures are formed in the cytoplasm during viral activation. The similar infection dynamics we observed in strain Ors24 to that of *C. reinhardtii* CC-2937 suggests that latent virus activation is taking place in both cultures during active growth. The phylogenetic proximity of the viruses involved, together with the previous discovery of a large clade of endogenous giant viruses in diverse green algae, indicates that a distinct viral lineage is associated with a range of different green algae in nature.

**Fig. 5.**
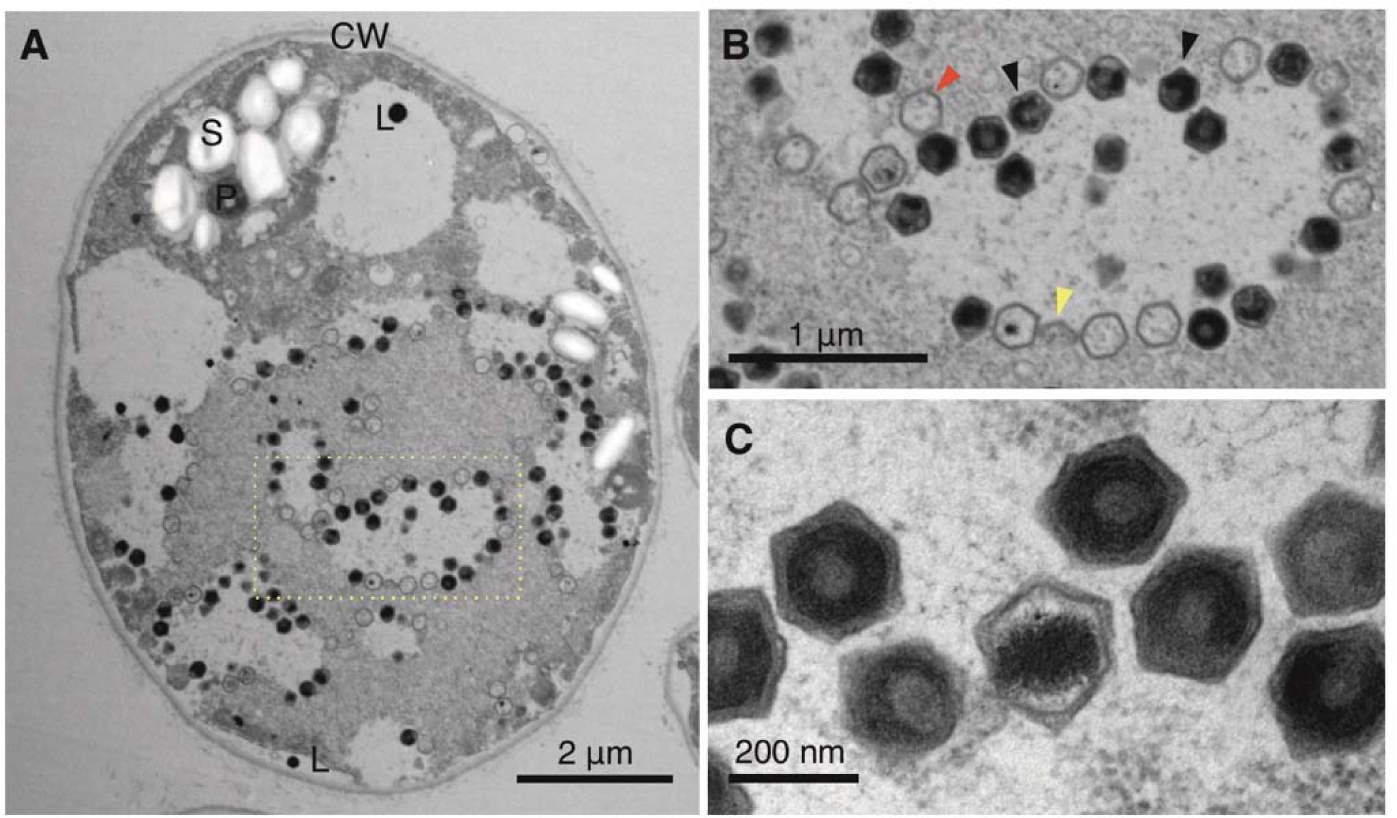
Transmission electron microscopy of ultra-thin sectioned Ors 24 *Chlamydomonas* sp. cells. **(A)** Infected *Chlamydomonas* sp. cell in early exponential phase (left). The nucleus and chloroplast are not clearly distinguishable. **(B)** Enlargement of the section delimited by a rectangle showing hexagonal viral particles. Virion production in a clearly delineated, lighter colored area with virions in later stages of completion accumulating at the edges of the production area, i.e. the virus factory/viroplasm. Red and black arrows indicate empty and full capsids, respectively. Yellow arrows indicate partially assembled capsids. **(C)** Enlarged picture of assembled virions. P – pyrenoid; CW – cell wall; L – lipid vesicle/plastoglobule; S – starch sheath.

## Conclusions

Punuivirus is the first virus known to infect *C. reinhardtii,* and its novel infection strategy opens multiple intriguing avenues of inquiry for future research. Firstly, it remains unclear how this virus can successfully integrate and excise its genome during different stages of infection, and given the size of its genome, it will be important for future work to examine the mechanistic details of this process and identify the enzymes involved. Secondly, it remains unclear what signals induce GEVE activation and virion production. Only a small fraction of cells produces virions, even during peak viral activity, suggesting that population heterogeneity during cellular growth plays a role or chemical signaling is involved. As virion production appears to peak during mid-exponential or stationary phase growth, we speculate that a buildup of metabolic byproducts may signal viral activation. Lastly, it is remarkable to consider that the GEVE genome encodes a variety of selfish genetic elements that are expressed, and in some cases can mobilize to other areas of the host and viral genomes. Among these, Fanzor elements are programmable RNA-guided nucleases that are of interest for genetic engineering applications, and in this context, one may consider that Punuivirus is a vector for these enzymes as part of its normal infection program. Undoubtedly, it will be revealing to understand the molecular details of how this process occurs during infection, as well as the long-term consequences of the multi-partite co-evolution between virus, host, and selfish genetic elements. Further research into this novel host-virus system is likely to uncover intriguing details that will be important for understanding viral infection strategies, as well as potentially informing biotechnology applications.

The latent infection program of Punuivirus is likely a common strategy among large protist viruses in nature that has traditionally been overlooked due to methodological challenges. For example, most cultivated giant viruses have been discovered due to their pronounced impact on cultures of their host (i.e. “culture crashes”), and the lack of any clear phenotypic effect of a latent virus, even during peak viral production, has likely impeded the earlier discovery of this phenomenon. Studies dating as far back as the 1970s observed viral production in otherwise healthy cultures of green algae and speculated that it may be due to the activity of latent viruses, but this was difficult to prove owing to technological limitations and the possibility of environmental contamination [12,13]. Our demonstration of GEVE activity in *C. reinhardtii* together with our discovery of widespread latent viruses in freshwater *Chlamydomonas* isolates, put these earlier observations into sharp focus and revives the view that latency is commonplace in large DNA viruses of protists.

A central aspect of Punuivirus latency is its ability to integrate into the *C. reinhardtii* genome, but viral integration is not necessarily a requirement for long-term persistent infections. Indeed, some virulent nucleocytoviruses can stably co-exist with their hosts by infecting only a subset of the population, thereby leading to long-term viral persistence without host population collapse [37,38]. Moreover, other giant viruses have low virulence in a particular host but appear to compensate with a broader host range [39]. We surmise that the integration of Punuivirus into the *C. reinhardtii* genome provides an added benefit to the virus by ensuring that it can be maintained even during long periods of host dormancy, for example in the durable zygospores that form during sexual reproduction. Zygospores are highly resistant to environmental perturbations, remaining viable in soil for several years [40]. Integration into the genomes of these cells may therefore allow Punuivirus to persist through seasonal transitions. Given the prevalence of GEVEs across eukaryotes [8–10], it is clear that genome integration is a common strategy among giant viruses. Indeed, recent studies have also reported endogenous viral elements derived from other lineages of eukaryotic DNA viruses, such as those of the recently discovered *Mirusvirocota* phylum [41]. Altogether, these findings suggest that latency and genome integration are employed by a broad group of large eukaryotic viruses during infection.

## Materials and Methods

### Maintenance of and culture conditions for C. reinhardtii CC-2937

*Chlamydomonas reinhardtii* strain CC-2937 was acquired from the Chlamydomonas Resource Center (Minneapolis, MN, USA) and maintained on 2% agar TAP media (#T8224, Plant Phytotech Labs, Lenexa, KS, USA) slants supplemented with 4 g L^-1^of yeast extract and 1 mL L^-1^ of glacial acetic acid, adjusted to pH 7 with glacial acetic acid. Liquid TAP media was prepared identically omitting the agar and yeast extract. All liquid and agar cultures were maintained at 24 °C under a 12:12-h light:dark cycle at an intensity of 100 μmol quanta m^-2^ s^-1^. Liquid cultures were agitated using an orbital shaker (Fisherbrand™ Multi-Platform shaker, Thermo Fisher Scientific, Waltham, MA, USA) at 150 rpm in conical flasks with a total capacity twice that of the media volume used. Cell densities were measured using the CytoFLEX-S Flow Cytometer (Beckman Coulter, Brea, CA, USA) equipped with violet (405 nm), and blue (488 nm) lasers. Chlorophyll autofluorescence was excited by the 488 nm blue laser and collected using a 780/60 nm band pass filter. A total of 10,000 events were recorded per measurement, with a medium flow rate (30 μL min^-1^). Cultures were diluted accordingly to maintain reads between 100 - 1,500 events μL^-1^. The Forward scatter and chlorophyll autofluorescence channels were set with automatic thresholds and gain values of 42 and 124, respectively.

### Genomic DNA extraction

High molecular weight genomic DNA (gDNA) was isolated from *C. reinhardtii* CC-2937 late-exponential cultures (∼10^7 cells mL^-1^). First, 50 mL of the culture was centrifuged at 4,500 g for 5 min in a Sorvall ST1R Plus-MD centrifuge with the TX-400 rotor (Thermo Fisher Scientific, Waltham, MA, USA). The resulting pellet was washed once with PBS 1X and gently resuspended in 5 mL of SDS buffer (50 mM Tris-HCl pH 8, 200 mM NaCl, 20 mM EDTA, 2% SDS) and 5 mL of CTAB buffer (100 mM Tris-HCl pH 8, 20 mM EDTA pH 8, 1.4 M NaCl, 2% CTAB, 1% PVP M.W. 40,000) preheated at 65 °C. Five μL of RNase A (100 mg mL^-1^) and Proteinase K (20 mg mL^-1^) were added and incubated at 65 °C for 1 h, mixing every 15 min. The lysate was centrifuged at 4,500 g for 5 min and decanted into a new falcon tube. One volume of phenol-chloroform (1:1) was added and mixed by inversion for 10 min, followed by centrifugation at 3,000 g for 10 min. The supernatant was transferred to a new tube using wide-bore pipette tips and extracted again using one volume of chloroform. Five μL of proteinase K (20 mg mL^-1^) and 10 μL of RNase A (100 mg mL^-1^) were added and incubated at 50 °C for 1 h, followed by the addition of one volume of chloroform and centrifugation at 3,000 g for 10 min. The DNA in the supernatant was precipitated with 2.5 volumes of cold 100% ethanol, and recovered by centrifugation at 4,500 g for 5 min. The pellet was transferred to a DNA LoBind tube containing 70% ethanol, and dried at 39 °C for approximately 10 min. Finally, the DNA was resuspended in 400 μL of molecular grade water and stored at 4°C.

### DNA shearing, cleanup and library preparation for long-read sequencing

The extracted gDNA was sheared 30 times with a 27 G syringe needle and purified using 0.7x AMPure XP beads (Beckman Coulter Inc., Indianapolis, IN, United States) for 15 min. The DNA was eluted in 60 μL of preheated elution buffer (10mM Tris-Cl pH8) for 20 min at 40 °C. The DNA purity was measured with a NanoDrop 2000 spectrophotometer (Thermo Fisher Scientific, Waltham, MA, USA), and integrity was assessed by performing agarose gel electrophoresis, and running 4200 TapeStation Genomic DNA ScreenTape assays (Agilent Technologies, Santa Clara, CA, USA). The DNA concentration was measured using a Qubit fluorometer, and approximately 3.5 μg of DNA was used for library preparation using the SQK-LSK114 Ligation Sequencing Kit V14 from Oxford Nanopore (Oxford Science Park, UK) with modifications. The formalin-fixed paraffin-embedded (FFPE) repair step was omitted, and the end-prep step was performed using the NEBNext Ultra II End Repair Module (New England BioLabs, Ipswich, MA) according to the manufacturer’s instructions. The end-prepped DNA was diluted with two volumes of elution buffer and cleaned up with 1x AMPure XP beads as described previously, using 60 μL of elution buffer. For the adapter ligation, the reaction was incubated for 1h, then diluted with 1 volume of elution buffer, and cleaned with 0.8x AMPure XP beads, which were washed twice with 250 μL of a mix of SFB:LFB buffer (1:2). The DNA was eluted with 30 μL of the provided elution buffer as described before. About 2.7 μg of library was recovered which was used to load a PromethION Flow Cell (R10.4.1) three times (370 ng per load) after washing it for 2 h every 24 h using the Nanopore EXP-WSH004 Flow Cell Wash Kit. Prior to this run, two sequencing attempts without flow cell reloads were performed using 1 μg of unsheared and sheared gDNA as the input for library preparation following the kit instructions.

### Genome assembly and polishing

The raw reads were base-called in real time using the MinKNOW software (v23.07.5) and Guppy (v7.0.9) with the high accuracy model (400bps, 5khz). Reads generated with the three sequencing runs were pooled, and those shorter than 1 kb were discarded. A draft genome assembly was generated with Flye v2.8.3 [42] with the options --nano-raw and a genome size of 120 Mb. The assembly was polished using long reads and four rounds of Racon v1.4.20 (https://github.com/isovic/racon) with default settings, followed by one round of Medaka v1.11.1 (https://github.com/nanoporetech/medaka) specifying the model r1041_e82_400bps_hac_v4.2.0. The medaka consensus assembly was further polished with two rounds of Racon using Illumina short reads that had previously been generated for the CC-2937 strain (NCBI SRA accession SRR1734616) [43]. The estimated genome size recovered was ∼110 Mb, which included 77 contigs and 2 scaffolds with an N50 value of 3.9 Mb. Contigs below 10 kbp were filtered for downstream analyses.

### GEVE contig identification

ViralRecall v2.1 [23] was run on the final polished assembly (using the contig screening parameter ‘-c’) to identify the contig(s) containing the NCLDV marker genes. One contig (contig_536) was found to contain clear signatures of NCLDV endogenization, an manual inspection confirmed that this was a GEVE flanked by regions that were homologous to chromosome 15 in the latest *C. reinhardtii* reference assembly (v6) [22]. BLASTn v2.12.0 [44] was used to search of the polished contig_536 against this chromosome (evalue 1e-20, length > 500 and percent identity >= 90) and synteny results were visualized using the R package gggenomes (https://thackl.github.io/gggenomes/). The TIRs were annotated using Minimap2 [45] by mapping the contig against itself as described previously [7]. The precise GEVE region was determined by the TIR boundaries, minimap2 alignment and manual comparison of the flanking sequence among CC-2937, the reference genome, and the genomes of two other field isolates (CC-1952, CC-2937 [26]). GEVE relics were identified by BLASTn searches using the GEVE TIRs as query sequences. The repeat content was calculated with Tandem Repeats Finder v4.09 [46] with parameters described elsewhere [47]. The Repeat and GC fractions were calculated from 10 kb non-overlapping sliding windows. Tetranucleotide frequency deviation of the GEVE compared to the rest of chromosome 15 was calculated using methods previously described [8].

### Functional annotation

Protein function of the GEVE predicted ORFs was retrieved from the full annotation table produced by ViralRecall. Additional annotations were obtained using eggNOG-mapper v2 [48] and HHpred for selected proteins [49]. Fanzor elements, including partial or variant configurations, were identified through a local BLASTn search in QIAGEN CLC Main Workbench v7.9.1 using default parameters. A Fanzor nuclease sequence alignment was generated using Muscle v5.1 [50] using default parameters. The alignment was trimmed using trimAl v1.4 [51] prior to tree generation removing positions with a gap in 90% or more of sequences. IQ-TREE v2.1.4 [52] was used to generate trees from the alignments, using ModelFinder Plus [53] for model selection and 1000 ultrafast bootstraps [54]. The best tree of 10 runs was selected.

### Exponential-to-stationary growth experiment and RNA sequencing of C. reinhardtii CC-2937

*Chlamydomonas reinhardtii* strain CC-2937 was taken from a slant and cultured in liquid TAP media. A starter culture was maintained at a cell density of approximately 150 - 200 × 10^3^ cells mL^-1^ through daily dilution. From this starter culture, new cultures were initiated each day for five consecutive days, starting from March 31, 2023, with an initial cell density of approximately 100 × 10^3^ cells mL^-1^. The cell density of each culture was monitored daily using flow cytometry as previously described. On the seventh day (April 6, 2023), 1.5 mL of sample from each culture were harvested by centrifugation (4,000 g, 4 min, 4°C). The supernatant was discarded, and the cell pellets, containing between 3-8 million cells each, were rapidly frozen in liquid nitrogen and stored at −80°C for future analysis.

RNA was extracted from cell pellets using TRIzol Plus RNA Purification Kit (Invitrogen, 12183555), following the manufacturer’s instructions. Total RNA was quantified using a Qubit™ RNA HS kit (Invitrogen, Q32852) and the quality was assessed with the HS RNA ScreenTape on an Agilent TapeStation system. RNA was converted into a strand-specific library using Illumina’s Stranded Total RNA Prep, Ligation with Ribo Zero Plus Sample Prep Kit (Illumina, 20040529) for subsequent cluster generation and sequencing on Illumina’s NovaSeq 6000. Supplemental Probes (table S3) were used in the Hybridize Probes step in the RiboZero Plus reactions. The libraries were enriched by 13 cycles of PCR, validated using Agilent TapeStation, and quantitated by qPCR (P5 Primer: AATGATACGGCGACCACCGA, P7 Primer: CAAGCAGAAGACGGCATACGAGAT). Individually indexed cDNA libraries were pooled and sequenced on NovaSeq 6000 SP 150 cycle PE using Illumina NovaSeq Control Software v1.8.0. The BCL files were converted to FASTQ files, and adapters were trimmed and demultiplexed using bcl2fastq Conversion Software.

### Gene prediction of the RNA-Seq assembled transcripts

A transcriptome co-assembly was generated from raw RNA-Seq illumina reads using rnaSPAdes v3.13.0 [55]. The polished *C. reinhardtii* CC-2937 assembly was soft-masked using tantan v22 [56], and then assembled transcripts were mapped onto this reference using BLAT v35 [57] (parameters “-minIdentity=92”) to obtain a psl file. The psl file was converted to a hints file using the blat2hints.pl PERL script provided with AUGUSTUS (https://github.com/nextgenusfs/augustus), and genes were then predicted using AUGUSTUS v. 3.5.0 [58] with the hints file and polished assembly used as input (parameters --species=chlamy2011 --softmasking=1). For the GEVE region, the coding density was lower than expected for a viral genome, likely because AUGUSTUS was not designed for viral gene prediction. To resolve this issue, AUGUSTUS gene predictions in the GEVE region were replaced with genes predicted using Prodigal v2.6.3 [59] (default parameters). To estimate the expression level of genes in the different RNA-Seq experiments, raw RNA-Seq reads were trimmed with Trim Galore v0.6.4 (https://github.com/FelixKrueger/TrimGalore, parameters “--length 36 -q 5 --stringency 1”) and then mapped onto the predicted transcripts using CoverM v0.4.0 with a minimum covered fraction of 20% (https://github.com/wwood/CoverM).

### Differential expression analysis

A differential expression analysis of the RNA-Seq count data was performed with the DESeq2 package [60]. The reference level for all contrasts was set to the youngest culture (3 days post-inoculation). Significant differentially expressed genes (DEGs) were identified based on an absolute log2-fold change (LFC) of 1.5 (Wald test: lfc threshold=0.585, alpha = 0.05) and an adjusted p-value < 0.05. The shrunken LFCs were estimated using the “normal” shrinkage estimator and visualized with volcano plots generated by the EnhancedVolcano (https://github.com/kevinblighe/EnhancedVolcano) R package. DEGs were clustered using self-organizing maps (SOMs) via the kohonen package in R [61]. We used the SOM codebook vectors in combination with K-means clustering to determine the optimal number of gene clusters. Then, we performed an unsupervised hierarchical clustering using the Ward method and Euclidean distance on the SOM codes to assign each gene to a cluster. The normalized DESeq2 counts belonging to the GEVE genes were scaled across samples with the min-max method prior to visualization on a heatmap generated using the pheatmap R package (https://github.com/raivokolde/pheatmap).

### Detection of viral particles by PCR and qPCR

A loopful of freshly growing CC-2937 cells on TAP agar plates (7-days old) was used to inoculate a 25 mL TAP media starter culture, which was grown for six days to a density of 9.5 × 10^6^ cells mL^-1^. The starter culture was then used to inoculate four flasks with 125 mL of TAP media to a final cell density of ∼3 × 10^5^ cells mL^-1^. For a total of 11 days, cell density was monitored daily with flow cytometry, and cells were observed with a Nikon Ti2-E inverted microscope with transmitted illumination. Five hundred µL samples were collected and centrifuged at 900 g for 1 min, and the supernatants were then filtered using 0.45 µm PES syringe filters and stored at 4 °C until further processing. Unpackaged host DNA contamination in the filtrates was removed by treating the samples with 1 U of DNAse I (#EN0521, Thermo Fisher Scientific, Waltham, MA, USA) per 8 µL of sample, following manufacturer’s instructions and stored at −20 °C until use. Viral DNA was detected by PCR using 40 cycles and 2 µL of the DNAse-treated filtrates as template in 10 µL final volume reactions, using the GEVE major capsid protein (*mcp*) primer pair, and conditions described in a previous study [21]. Host DNA contamination was detected by amplifying the ITS1-5.8S-ITS2 region using the primer pair Fw_ITS1/Rv_ITS4 [62,63] and the same conditions used previously.

To quantify free virions in terms of MCP copies µL^-1^ using qPCR, we initially generated a standard curve constructed from a dilution series ranging from 2.82E+00 to 1.41E+05 molecules per µL of a linear construct (Twist Bioscience, South San Francisco, CA, USA) containing a fragment of the GEVE *mcp* gene (GEVE_395) (5’ CAATCCGCCCTCACTACAACCGGAGTGCTACTATTTCAGCCAGCTCACAAACCAGAAGAACGGGTCCATTACCATAGGCAACTTGGATGCTTCGATGTACCTGGATTACGTGTATCTGGACACAGATGAGCGCAAGAAGTTTGCCCAAGCCGCTCACGAATACCTGGTGGAGCAGCTGCAGTATACCGGCGAGGAGTCGCTGCAGGGAAGCCAGGGCAAGGTGAAGCTGAGCCTGAACCACCCCGTTAAGGAGCTGATTTGGGTGATGCAGAAGGATGACTGGCTGACCAACACCGGCGCCAGGGTGATTGTGCCTACCTCTGCTACTCTGGCGTCGATGAGGGA 3’). The target sequence was amplified with the primers GEVE_MCP_qPCR forward (5□ GCAAGAAGTTTGCCCAAGCCGC 3□) and reverse (5□ CTCAGCTTCACCTTGCCCTGGC 3□) which amplified a product of 100 bp. We conducted triplicate qPCR reactions of 24 μL containing 1x Platinum™ SYBR™ Green qPCR SuperMix-UDG w/ROX (#11744500, Thermo Fisher Scientific, Waltham, MA, USA), a final concentration of 300 nM of each primer, and 4 µL of the DNAse-treated filtrates. The thermal cycling was performed in the CFX96 Real-Time PCR system (BioRad, Hercules, CA, USA) using the following settings: 50 °C for 2 min, 95 °C for 2 min, 45 cycles of 95 °C for 15 s followed by 60 °C for 30 s, and a final melting curve from 65 °C to 95 °C. Between all qPCR runs, the equation of the standard curve was Cq = 46.09 - 3.29 log10 (*mcp* copies mL^-1^) and the R2 value was 0.94.

### Viral population detection through flow cytometry

After 15 days of incubation, and considering the low viral loads, the contents of the four replicate flasks used for qPCR were pooled to ensure sufficient material for concentration via Tangential Flow Filtration (TFF) before detecting viral particles by flow cytometry. Cultures were centrifuged at 4,695 g, for 20 min at 4°C to pellet the cells in a Sorvall ST1R Plus-MD centrifuge with the TX-400 rotor (Thermo Fisher Scientific, Waltham, MA, USA). The collected supernatant was filtered sequentially through decreasing pore size filters using a peristaltic pump. A total of six 5 µm filters, two 3 µm filters, and one 0.8 µm filter were used to filter out cellular debris without clogging. About 350 mL of filtrate was recovered and stored overnight at 4°C. Viral particles concentration was performed using a 100 kDa (MWCO) PES Vivaflow 200 TFF unit (Sartorius, Göttingen, Germany), keeping an inlet pressure <10 psi, to a final volume of approximately 25 mL (∼14-fold concentration). Fifteen mL of this sample was further concentrated with an Amicon 100 kDa Ultra Centrifugal Filter (Millipore Sigma, St. Louis, MO) to about 1.1 mL (total ∼190-fold concentration). This concentrate was diluted 1:2 with molecular grade water and 50 µL was fixed with 25% glutaraldehyde (stored at 4°C, Electron Microscopy Sciences, Hatfield, PA, USA) to a final concentration of 0.25%, for 20 min in the dark at room temperature. The rest of the sample was filtered using a 0.8 µm PES filter, and a 50 µL aliquot was fixed as described before. This step was repeated with a 0.45 µm PES filter to visualize the particles using different pore size cutoffs (fig. S3D-F).

Based on the virus detection and enumeration protocol by Brussaard et al. [64,65], the glutaraldehyde fixed samples (2x diluted) were further diluted (50x) with TE-buffer (10 mM Tris-HCl pH 7, 1mM EDTA pH8) and stained with nucleic acid-specific SYBR Safe (diluted to 1X in the TE-buffer, Thermo Fisher Scientific, Waltham, MA, USA) for 20 min at 80 °C. Particles with virus-like green fluorescence were excited using the 405 nm violet and 488 nm blue lasers using the CytoFLEX-S flow cytometer. The Violet Side Scatter (V-SSC, 200 gain), and the Green Fluorescence (B525, 2,000 gain) channels were fitted with a 405/10 and a 525/40 band pass filter, respectively [66,67]. The thresholds were set to 1200 for V-SSC, and 20,000 for B525 to keep the abort rate below 1 %.

As a positive control for the detection of large DNA viruses, we used Paramecium bursaria chlorella virus 1 (PBCV-1) and Acanthamoeba polyphaga mimivirus lysates. For the PBCV-1 positive control, the B525 channel threshold was decreased to 15,000, since it had a slightly lower green fluorescence signal under the set parameters. Both controls were easily detected (fig. S3A-B). The filtrated (permeate; < 100 kDa) collected from the Vivaflow concentration unit was treated identically and used as a negative control (fig. S3I).

### Identification of virions using proteomics

Three replicates were conducted by filling two 1 L Erlenmeyer flasks with 500 mL of TAP media and inoculating each with a loopful of freshly growing CC-2937 cells from 7-day-old TAP agar plates. The flasks were incubated for 9 days, and the cultures were concentrated using a 100 kDa PES Vivaflow 200 TFF unit, following previously described methods, to a final volume of approximately 22 mL (∼40-fold concentration). The concentrated supernatant (21 mL) was distributed into 1.5 mL tubes and centrifuged at 15,000 g for 1 hour at 4°C. From each tube, 1400 µL of the supernatant was carefully removed, and the remaining volumes were pooled into a single tube, which was then centrifuged under the same conditions for an additional hour. The supernatant was discarded, and the resulting pellets were frozen at −80°C until proteome analysis.

Protein was solubilized in S-trap lysis buffer (10% w/v SDS in 100 mM triethylammonium bicarbonate pH 8.5). Proteins were reduced using DTT (4.5 mM) and alkylated with IAA (10 mM). Unreacted IAA was quenched with DTT (10 mM) and samples were acidified using o-phosphoric acid. Protein was precipitated using methanol and incubation at −80°C overnight. An aliquot of each sample corresponding to 100 μg protein was loaded onto a mini S-trap (Protifi, Fairport, NY, USA) and washed with methanol. Proteins were then digested overnight with 2 μg trypsin in 25 μl 50 mM triethyammonium bicarbonate (pH 8.5).

LC-MS/MS was performed in duplicate using a Thermo Fisher Scientific Vanquish Neo HPLC and autosampler (Waltham, MA, USA) system controlled by Chromeleon 7.2.10 coupled online to a Bruker timsTOF fleX mass spectrometer via a Bruker Captive Spray ion source (Billerica, MA, USA). Three micrograms (3 μl) peptide solution were separated on a PharmaFluidics 50 cm μPAC capLC C18 column (Thermo Fisher Scientific, Waltham, MA, USA) at a flow rate of 350 nl min^-1^ in an oven compartment heated to 40°C. The LC gradient used started with a linear increase (solvent A: 2% acetonitrile, 98% water and 0.1% formic acid; solvent B: 80% acetonitrile, 20% water and 0.1% formic acid) from 2% B to 10% B over 3 min, followed by a linear increase from 10% B to 50% B over 88 min followed by a wash of 4 minutes at 98% B.

For the DDA-PASEF acquisition mode, 1 survey TIMS-MS and 10 PASEF MS/MS scans were performed per acquisition cycle. We analyzed an IM range from 1/K0 = 0.6 to 1.6 V-s/cm2 using equal ion accumulation and ramp time in the dual-TIMS analyzer of 100 ms each. Suitable precursor ions for MS/MS analysis were isolated in a window of 2 Th for m/z < 700 and 3 Th for m/z > 700 by rapidly switching the quadrupole position in sync with the elution of precursors from the TIMS device. The collision energy was lowered stepwise as a function of increasing IM, starting from 20 eV for 1/K0 = 0.6 Vs/cm2 and 59 eV for 1/K0 = 1.6 Vs/cm2 making use of the m/z and IM information to exclude singly charged precursor ions with a polygon filter mask and further used “dynamic exclusion” to avoid resequencing of precursors that reached a “target value” of 20,000 au. The IM dimension was calibrated linearly using three ions from the Agilent ESI LC/MS tuning mix (m/z, 1/K0: 622.0289, 0.9848 Vs/cm2; 922.0097, 1.1895 Vs/cm2; and 1221.9906, 1.3820 Vs/cm2).

Data files were processed with Mascot Distiller 2.8.5 (Matrix Science, Boston, MA) using the default settings for data generated using Bruker timsTOF instruments. Processed data were then searched using Mascot 2.8.3 (Matrix Science, Boston, MA, USA). The search utilized the UniProt reference *Chlamydomonas reinhardtii* proteome database, a common protein contaminant database and the FASTA formatted GEVE proteome. The search assumed trypsin-specific peptides with the possibility of 2 missed cleavages, a precursor mass tolerance of 100 ppm and a fragment mass tolerance of 0.1 Da, a fixed modification of carbamidomethyl at Cys and the variable modifications of oxidation of Met and cyclization of a peptide N-terminal Gln to pyro-Glu. To identify a final list of proteins packaged in the virions, only proteins that were detected at least twice across all technical runs, and with a Mascot database search score >=35 were retained. Additionally, proteins with a lower score were also retained if at least two Peptide Spectrum Match (PSM) were recorded in at least two runs.

### Electron microscopy of C. reinhardtii CC-2937 concentrated supernatants

The remaining concentrated supernatant (∼5.5 mL) was used for viral particle screening with transmission electron microscopy. Viral particles were pelleted by centrifugation at 15,000 g for 1 hour at 4°C. Approximately 1300 µL of the supernatant was removed, and the remaining volume was pooled into a single tube and topped up to 1.5 mL with molecular-grade water. After homogenization, the sample was filtered through a 0.8 µm PES filter (13 mm diameter) and centrifuged for an additional hour under the same conditions. The supernatant was discarded, and the pellet was resuspended in 100 µL of molecular-grade water, then filtered through a 0.45 µm PVDF filter (4 mm diameter). A non-GEVE strain supernatant (*C. reinhardtii* CC-2935) was also concentrated and pelleted in the same manner, serving as a negative control to differentiate VLPs from non-VLPs. Flow cytometry was performed on these samples, further confirming that concentrating the non-GEVE strain CC-2935 results in the absence of a small particle population compared to CC-2937 (fig. S3G-H).

Formvar/Carbon 200-mesh copper grids (Electron Microscopy Sciences, Hatfield, PA) were hydrophilized with UV-C light (254 nm) radiation for 2h in a PCR Station Enclosure (#3970305, Labconco, Kansas City, MO, USA). Ten µL of the concentrated sample were placed onto the grid and incubated for 10 min. The excess sample was blotted with filter paper and stained with 3 µL of uranyl acetate 2 % for 30 seconds. This step was repeated once, and the grid was allowed to dry at room temperature. The VLPs were visualized with a JEM 2100 Transmission Electron Microscope (JEOL, Tokyo, Japan) operated at 200 kV. The average diameter of 23 viral particles (4 measurements per virion) was measured with the software ImageJ [68].

### Assessing the prevalence of viral induction in freshwater Chlamydomonas sp. field isolates

Water samples were collected in 2016 from Örsjön (56.2828485, 14.6838229) and Krageholmssjön (55.50159, 13.74462), two lakes located in southern Sweden, using a 10 μm plankton net. Single *Chlamydomonas*-like green algal cells were isolated by hand using an inverted microscope. Individual cells were washed with five drops of filtered lake water (0.2 µm pore size) and placed into 96-well plates containing 0.5X MWC+Se media diluted with filtered lake water [69]. Once wells turned visibly green, the cultures were sequentially transferred to larger wells, until finally transferring them to 30 mL of 1X MWC+Se media in Nunc T25 tissue culture flasks (#169900, Thermo Fisher Scientific, Sweden). A final total of 18 and 20 monoalgal cultures were established from Örsjön and Krageholmssjön, respectively. These monocultures were maintained at 15°C using a 12:12 h light:dark cycle and reduced light intensity (10 μmol quanta m^-2^ s^-1^) with monthly transfers to fresh media. For the experiments and DNA extraction growth was increased by elevating light intensity to 90 μmol quanta m^-2^ s^-1^.

To confirm that isolates were *Chlamydomonas* sp., we sequenced the 18S rRNA gene following methods described elsewhere [70]. For phylogenetic analysis we aligned the 18S sequences of two representative isolates from both lakes (Ors24 and Kgh138), the *C. reinhardtii* CC-2937 strain, and a collection of reference 18S sequences from the PR2 database [71]. To select references, we compared the 18S sequences of the isolates to the PR2 sequences using BLASTn, and retained the top 100 hits for each. We then dereplicated the reference 18S sequences at 99% identity using CD-HIT [72] and aligned all sequences together using Muscle v5.1 [50]. We trimmed the alignments to remove all sites with >20% gaps using trimAl v1.4 [51], and constructed the phylogeny using IQ-TREE v2.03 [52] with the TIM2+F+I+G4 model, as chosen by ModelFinder [53], and 1000 ultrafast bootstraps.

To screen the monocultures for endogenous viruses, we grew them to mid-exponential phase (approximately 75 × 10^3^cells mL^-1^), followed by DNA extraction and PCR amplification of the viral *mcp* gene using the degenerate primers, mcp Fwd and mcp Rev, which target the *mcp* sequence of large algal viruses [73]. For DNA extraction, cultures were centrifuged at 3,000 g for 10 minutes, and after most of the supernatant was decanted, the pellet was gently resuspended in the remaining liquid. The cell fraction was further transferred to 1.5 mL tubes, centrifuged, and the resulting pellets were stored at −80 °C until further processing. DNA was extracted from the cell pellets using the DNeasy Plant Mini kit (Qiagen, Valencia, CA, USA) with modifications. First, the pellets were transferred to 2 mL screw-cap tubes with a small amount of glass beads (212-300 μm) and shock frozen at −150°C for 5 min. Then, 100 μL of buffer AP1 was added, and lysis was performed with the TissueLyser II (Qiagen, Valencia, CA) at 30 Hz, for 30-60 s. An additional 300 μL of buffer AP1 was added, and the procedure was continued according to the manufacturer’s instructions. The DNA was quantified using a NanoDrop 2000 spectrophotometer (Thermo Fisher Scientific, Waltham, MA, USA). The PCR reactions were performed in a final volume of 25 μL with final concentrations of 1.5 mM MgCl2, 0.25 mM dNTPs, 0.2 mg mL^-1^ BSA, 0.8 μM of primers, 0.12 U of AmpliTaq (Thermo Fisher Scientific, Waltham, MA, USA), and approximately 40 ng of DNA. Initial denaturation was performed at 94°C for 4 min, followed by 39 cycles of denaturation at 94 °C for 30 s, annealing at 50 °C for 45 s, elongation at 72 °C for 1 min, and a final elongation at 72 °C for 10 min.

The two representative isolates from both lakes (Ors24 and Kgh138) were selected for purification and sequencing of the *mcp* PCR products using the MinElute PCR Purification Kit (Qiagen, Valencia, CA, USA), following the manufacturer’s instructions. Sequencing was performed in an Applied Biosystems 8-capillary 3500 Genetic Analyzer using the BigDye Terminator Cycle Sequencing Kit (Thermo Fisher Scientific, Waltham, MA, USA). High-quality chromatograms were manually trimmed and aligned in Geneious® 11.0.2, using the Geneious alignment algorithm. An online BLASTn search of the consensus sequence revealed matches with other viral MCP sequences of large dsDNA viruses. For phylogenetic analysis, we aligned the Ors24, Kgh138, and Punuivirus MCP sequences together with a set of homologous protein sequences from known green algal GEVEs and reference nucleocytoviruses available in the Giant Virus Database (https://faylward.github.io/GVDB/) using Muscle5 v5.1 (default parameters). The tree was constructed using IQ-TREE v2.03 using the -alrt support option and the LG+G+R10 substitution model. For the Ors24 strain we also obtained low-coverage long-read sequencing using the PacBio Revio Technology Platform with the multiplex HiFi protocol at the Uppsala Genome Center, Science for Life Laboratory. This sequencing recovered a near full-length viral family B polymerase, which we placed into a tree together with other representative viral sequences using the same methods as for the MCP tree.

To screen for VLPs inclusions using transmission electron microscopy, Ors24 was harvested at different stages of growth, including early exponential, mid-exponential, late exponential, and stationary phase (∼2, 4, 6 and 8 × 10^5^ cells mL^-1^, respectively). Sample preparation was performed according to Hoops & Witman [74] with modifications. Cells were double fixed with 4% glutaraldehyde (GA) in MWC+Se media for 15 min at room temperature and then transferred to 4% GA in 100 mM sodium cacodylate (NaCac) for 4 h at room temperature. The GA+NaCac was removed and 100 mM of NaCac buffer was added in the dark at 4°C. Samples were post-fixed in 1% osmium tetroxide water for 2 h at 4°C, and the pellets were dehydrated in graded ethanol series and embedded in epoxy resin (Agar 100) via acetone. Semi thin-sections (1.5 μm) were made using a Leica EM UC7 ultramicrotome with a glass knife and stained with Richardson’s solution [75] to examine the orientation of the tissue in the trimmed block. Ultra-thin sections (50 nm) were made using a Leica EM UC7 ultratome with a diamond knife. The sections were mounted on pioloform coated, single slot, copper grids and stained with uranyl acetate (2%, 30 min) and lead citrate (4 min). The grids were visualized using a JEOL JEM 1400 Plus transmission electron microscope (Jeol, Tokyo, Japan). A total of 40 icosahedral VLPs were measured for size, with the diameter of each VLP calculated as the average of three measurements across vertices.

## Acknowledgements

Sequencing was performed at the Genomics Sequencing Center which is part of the Fralin Life Science Institute at Virginia Tech. We thank Rich Helm for assistance with the proteomics and Stephen McCartney and Hongyu Wang for assistance with Transmission Electron Microscopy at the Nanoscale Characterization and Fabrication Lab, Virginia Tech. We are grateful to Chantal Abergel and James Van Etten for providing cultures of mimivirus and Paramecium bursaria chlorella virus 1, respectively, for their use as positive controls in our flow cytometry assays. The authors would also like to acknowledge support of the National Genomics Infrastructure (NGI) / Uppsala Genome Center and UPPMAX for providing assistance in massive parallel sequencing and computational infrastructure. We acknowledge the Microscopy and DNA Sequencing Facility at the Department of Biology, Lund University for assistance with Transmission Electron Microscopy and Sanger sequencing of the freshwater *Chlamydomonas* strains.

## Author contributions

Conceptualization: K.R., C.P.D.B., M.M., F.O.A.; Methodology: M.P.E.G., U.S., Z.K.B., R.J.C., P.W., A.M.J., W.K.R., M.S.C., C.K.C., K.R., C.P.D.B.; Investigation: M.P.E.G., U.S., Z.K.B., R.J.C., P.W., A.M.J., W.K.R., M.S.C.; Visualization: M.P.E.G., U.S., Z.K.B., R.J.C., P.W., F.O.A.; Supervision: K.R., C.P.D.B., F.O.A.; Provision of reagents: K.R., C.P.D.B., F.O.A.; Funding acquisition: K.R., C.P.D.B., F.O.A.; Writing – original draft: M.P.E.G., U.S., Z.K.B., F.O.A.; Writing – review and editing: M.P.E.G., U.S., Z.K.B., R.J.C., P.W., A.M.J., W.K.R., M.S.C., C.K.C., K.R., C.P.D.B., M.M., F.O.A.

## Data and materials availability

The raw data for the three long read sequencing runs, and the raw RNA-Seq reads have been deposited in the NCBI Short-Read Archive under the BioProject number PRJNA1131777. The assembled *C. reinhardtii* CC-2937 contigs, gene predictions, protein predictions, and alignments for the phylogenetic trees are available in Zenodo (DOI: 10.5281/zenodo.13645515).

## References

1. Feschotte C, Gilbert C. Endogenous viruses: insights into viral evolution and impact on host biology. Nat Rev Genet. 2012;13: 283–296.

2. Katzourakis A, Gifford RJ. Endogenous Viral Elements in Animal Genomes. PLoS Genet. 2010;6: e1001191.

3. Takahashi H, Fukuhara T, Kitazawa H, Kormelink R. Virus Latency and the Impact on Plants. Front Microbiol. 2019;10: 2764.

4. Horie M, Honda T, Suzuki Y, Kobayashi Y, Daito T, Oshida T, et al. Endogenous non-retroviral RNA virus elements in mammalian genomes. Nature. 2010;463: 84–87.

5. Holmes EC. The Evolution of Endogenous Viral Elements. Cell Host Microbe. 2011;10: 368.

6. Chiba S, Kondo H, Tani A, Saisho D, Sakamoto W, Kanematsu S, et al. Widespread endogenization of genome sequences of non-retroviral RNA viruses into plant genomes. PLoS Pathog. 2011;7: e1002146.

7. Bellas C, Hackl T, Plakolb M-S, Koslová A, Fischer MG, Sommaruga R. Large-scale invasion of unicellular eukaryotic genomes by integrating DNA viruses. Proc Natl Acad Sci U S A. 2023;120: e2300465120.

8. Moniruzzaman M, Weinheimer AR, Martinez-Gutierrez CA, Aylward FO. Widespread endogenization of giant viruses shapes genomes of green algae. Nature. 2020;588: 141–145.

9. Zhao H, Zhang R, Wu J, Meng L, Okazaki Y, Hikida H, et al. A 1.5-Mb continuous endogenous viral region in the arbuscular mycorrhizal fungus. Virus Evol. 2023;9: vead064.

10. Sarre LA, Kim IV, Ovchinnikov V, Olivetta M, Suga H, Dudin O, et al. DNA methylation enables recurrent endogenization of giant viruses in an animal relative. bioRxiv. 2024. p. 2024.01.08.574619. doi:10.1101/2024.01.08.574619

11. Gong Z, Zhang Y, Han G-Z. Molecular fossils reveal ancient associations of dsDNA viruses with several phyla of fungi. Virus Evol. 2020;6: veaa008.

12. Dodds JA. Viruses of marine algae. Experientia. 1979;35: 440–442.

13. Reisser W. Viruses and Virus-Like Particles of Freshwater and Marine Eukaryotic Algae — a Review. Archiv für Protistenkunde. 1993;143: 257–265.

14. Delaroque N, Maier I, Knippers R, Müller DG. Persistent virus integration into the genome of its algal host, Ectocarpus siliculosus (Phaeophyceae). J Gen Virol. 1999;80: 1367–1370.

15. Müller DG, Kawai H, Stache B, Lanka S. A Virus Infection in the Marine Brown Alga Ectocarpus siliculosus (Phaeophyceae). Bot Acta. 1990;103: 72–82.

16. Cock JM, Sterck L, Rouzé P, Scornet D, Allen AE, Amoutzias G, et al. The Ectocarpus genome and the independent evolution of multicellularity in brown algae. Nature. 2010;465: 617–621.

17. Gyaltshen Y, Rozenberg A, Paasch A, Burns JA, Warring S, Larson RT, et al. Long-Read-Based Genome Assembly Reveals Numerous Endogenous Viral Elements in the Green Algal Bacterivore Cymbomonas tetramitiformis. Genome Biol Evol. 2023;15. doi:10.1093/gbe/evad194

18. Filée J. Multiple occurrences of giant virus core genes acquired by eukaryotic genomes: the visible part of the iceberg? Virology. 2014;466–467. doi:10.1016/j.virol.2014.06.004

19. Salomé PA, Merchant SS. A Series of Fortunate Events: Introducing Chlamydomonas as a Reference Organism. Plant Cell. 2019;31: 1682.

20. Sasso S, Stibor H, Mittag M, Grossman AR. The Natural History of Model Organisms: From molecular manipulation of domesticated Chlamydomonas reinhardtii to survival in nature. 2018 [cited 24 Jun 2024]. doi:10.7554/eLife.39233

21. Moniruzzaman M, Erazo-Garcia MP, Aylward FO. Endogenous giant viruses contribute to intraspecies genomic variability in the model green alga Chlamydomonas reinhardtii. Virus Evol. 2022;8: veac102.

22. Craig RJ, Gallaher SD, Shu S, Salomé PA, Jenkins JW, Blaby-Haas CE, et al. The Chlamydomonas Genome Project, version 6: Reference assemblies for mating-type plus and minus strains reveal extensive structural mutation in the laboratory. Plant Cell. 2023;35: 644.

23. Aylward FO, Moniruzzaman M. ViralRecall-A Flexible Command-Line Tool for the Detection of Giant Virus Signatures in ’Omic Data. Viruses. 2021;13. doi:10.3390/v13020150

24. Aylward FO, Moniruzzaman M, Ha AD, Koonin EV. A phylogenomic framework for charting the diversity and evolution of giant viruses. PLoS Biol. 2021;19: e3001430.

25. Merchant SS, Prochnik SE, Vallon O, Harris EH, Karpowicz SJ, Witman GB, et al. The Chlamydomonas genome reveals the evolution of key animal and plant functions. Science. 2007;318: 245–250.

26. Lopez-Cortegano E, Craig RJ, Chebib J, Balogun EJ, Keightley PD. Rates and spectra of de novo structural mutation in Chlamydomonas reinhardtii. Genome Res. 2022; gr.276957.122.

27. Craig NL, Craigie R, Gellert M, Lambowitz AM. Mobile DNA II. 2002.

28. Kojima KK, Bao W. IS481EU Shows a New Connection between Eukaryotic and Prokaryotic DNA Transposons. Biology. 2023;12. doi:10.3390/biology12030365

29. Kawato S, Nozaki R, Kondo H, Hirono I. Integrase-associated niche differentiation of endogenous large DNA viruses in crustaceans. Microbiol Spectr. 2024;12: e0055923.

30. Brussaard CPD. Optimization of Procedures for Counting Viruses by Flow Cytometry. Appl Environ Microbiol. 2004 [cited 13 Aug 2024]. doi:10.1128/AEM.70.3.1506-1513.2004

31. Yoon PH, Skopintsev P, Shi H, Chen L, Adler BA, Al-Shimary M, et al. Eukaryotic RNA-guided endonucleases evolved from a unique clade of bacterial enzymes. Nucleic Acids Res. 2023;51: 12414–12427.

32. Dunigan DD, Cerny RL, Bauman AT, Roach JC, Lane LC, Agarkova IV, et al. Paramecium bursaria chlorella virus 1 proteome reveals novel architectural and regulatory features of a giant virus. J Virol. 2012;86: 8821–8834.

33. Gann ER, Xian Y, Abraham PE, Hettich RL, Reynolds TB, Xiao C, et al. Structural and Proteomic Studies of the Aureococcus anophagefferens Virus Demonstrate a Global Distribution of Virus-Encoded Carbohydrate Processing. Front Microbiol. 2020;11. doi:10.3389/fmicb.2020.02047

34. Fischer MG, Kelly I, Foster LJ, Suttle CA. The virion of Cafeteria roenbergensis virus (CroV) contains a complex suite of proteins for transcription and DNA repair. Virology. 2014;466–467: 82–94.

35. Birkholz EA, Morgan CJ, Laughlin TG, Lau RK, Prichard A, Rangarajan S, et al. An intron endonuclease facilitates interference competition between coinfecting viruses. Science. 2024;385: 105–112.

36. Goodrich-Blair H, Shub DA. Beyond homing: competition between intron endonucleases confers a selective advantage on flanking genetic markers. Cell. 1996;84: 211–221.

37. Yau S, Krasovec M, Benites LF, Rombauts S, Groussin M, Vancaester E, et al. Virus-host coexistence in phytoplankton through the genomic lens. Sci Adv. 2020;6: eaay2587.

38. Joffe N, Kuhlisch C, Schleyer G, Ahlers NS, Shemi A, Vardi A. Cell-to-cell heterogeneity drives host–virus coexistence in a bloom-forming alga. ISME J. 2024;18: wrae038.

39. Blanc-Mathieu R, Dahle H, Hofgaard A, Brandt D, Ban H, Kalinowski J, et al. A persistent giant algal virus, with a unique morphology, encodes an unprecedented number of genes involved in energy metabolism. J Virol. 2021;95. doi:10.1128/JVI.02446-20

40. Harris EH. Chlamydomonas as a model organism. Annu Rev Plant Physiol Plant Mol Biol. 2001;52: 363–406.

41. Collier JL, Rest JS, Gallot-Lavallée L, Lavington E, Kuo A, Jenkins J, et al. The protist Aurantiochytrium has universal subtelomeric rDNAs and is a host for mirusviruses. Curr Biol. 2023;33. doi:10.1016/j.cub.2023.10.009

42. Kolmogorov M, Yuan J, Lin Y, Pevzner PA. Assembly of long, error-prone reads using repeat graphs. Nat Biotechnol. 2019;37: 540–546.

43. Flowers JM, Hazzouri KM, Pham GM, Rosas U, Bahmani T, Khraiwesh B, et al. Whole-Genome Resequencing Reveals Extensive Natural Variation in the Model Green Alga Chlamydomonas reinhardtii. Plant Cell. 2015;27: 2353–2369.

44. Camacho C, Coulouris G, Avagyan V, Ma N, Papadopoulos J, Bealer K, et al. BLAST+: architecture and applications. BMC Bioinformatics. 2009;10: 421.

45. Li H. Minimap2: pairwise alignment for nucleotide sequences. Bioinformatics. 2018;34: 3094–3100.

46. Benson G. Tandem repeats finder: a program to analyze DNA sequences. Nucleic Acids Res. 1999;27: 573–580.

47. Payne ZL, Penny GM, Turner TN, Dutcher SK. A gap-free genome assembly of Chlamydomonas reinhardtii and detection of translocations induced by CRISPR-mediated mutagenesis. Plant Communications. 2023;4. doi:10.1016/j.xplc.2022.100493

48. Cantalapiedra CP, Hernández-Plaza A, Letunic I, Bork P, Huerta-Cepas J. eggNOG-mapper v2: Functional Annotation, Orthology Assignments, and Domain Prediction at the Metagenomic Scale. Mol Biol Evol. 2021;38: 5825–5829.

49. Zimmermann L, Stephens A, Nam S-Z, Rau D, Kübler J, Lozajic M, et al. A Completely Reimplemented MPI Bioinformatics Toolkit with a New HHpred Server at its Core. J Mol Biol. 2018;430: 2237–2243.

50. Edgar RC. Muscle5: High-accuracy alignment ensembles enable unbiased assessments of sequence homology and phylogeny. Nat Commun. 2022;13: 1–9.

51. Capella-Gutiérrez S, Silla-Martínez JM, Gabaldón T. trimAl: a tool for automated alignment trimming in large-scale phylogenetic analyses. Bioinformatics. 2009;25: 1972–1973.

52. Nguyen L-T, Schmidt HA, von Haeseler A, Minh BQ. IQ-TREE: A Fast and Effective Stochastic Algorithm for Estimating Maximum-Likelihood Phylogenies. Mol Biol Evol. 2014;32: 268–274.

53. Kalyaanamoorthy S, Minh BQ, Wong TKF, von Haeseler A, Jermiin LS. ModelFinder: Fast Model Selection for Accurate Phylogenetic Estimates. Nat Methods. 2017;14: 587.

54. Hoang DT, Chernomor O, von Haeseler A, Minh BQ, Vinh LS. UFBoot2: Improving the Ultrafast Bootstrap Approximation. Mol Biol Evol. 2017;35: 518–522.

55. Bushmanova E, Antipov D, Lapidus A, Prjibelski AD. rnaSPAdes: a de novo transcriptome assembler and its application to RNA-Seq data. Gigascience. 2019;8. doi:10.1093/gigascience/giz100

56. Frith MC. A new repeat-masking method enables specific detection of homologous sequences. Nucleic Acids Res. 2011;39: e23.

57. Kent WJ. BLAT--the BLAST-like alignment tool. Genome Res. 2002;12: 656–664.

58. Keller O, Kollmar M, Stanke M, Waack S. A novel hybrid gene prediction method employing protein multiple sequence alignments. Bioinformatics. 2011;27: 757–763.

59. Hyatt D, Chen G-L, Locascio PF, Land ML, Larimer FW, Hauser LJ. Prodigal: prokaryotic gene recognition and translation initiation site identification. BMC Bioinformatics. 2010;11: 119.

60. Love MI, Huber W, Anders S. Moderated estimation of fold change and dispersion for RNA-seq data with DESeq2. Genome Biol. 2014;15: 550.

61. Wehrens R, Buydens LMC. Self- and Super-organizing Maps in R: The kohonen Package. J Stat Softw. 2007;21: 1–19.

62. Hadi SIIA, Santana H, Brunale PPM, Gomes TG, Oliveira MD, Matthiensen A, et al. DNA Barcoding Green Microalgae Isolated from Neotropical Inland Waters. PLoS One. 2016;11: e0149284.

63. White TJ, Bruns T, Lee S, Taylor J, Others. Amplification and direct sequencing of fungal ribosomal RNA genes for phylogenetics. PCR protocols: a guide to methods and applications. 1990;18: 315–322.

64. Brussaard CPD, Payet JP, Winter C, Weinbauer MG. Quantification of aquatic viruses by flow cytometry. Manual of aquatic viral ecology. 2010;11: 102–107.

65. Brussaard CPD, Marie D, Bratbak G. Flow cytometric detection of viruses. J Virol Methods. 2000;85: 175–182.

66. Zucker RM, Ortenzio JNR, Boyes WK. Characterization, detection, and counting of metal nanoparticles using flow cytometry. Cytometry A. 2016;89: 169–183.

67. Zhao Y, Zhao Y, Zheng S, Zhao L, Zhang W, Xiao T, et al. Enhanced resolution of marine viruses with violet side scatter. Cytometry A. 2023;103: 260–268.

68. Schneider CA, Rasband WS, Eliceiri KW. NIH Image to ImageJ: 25 years of image analysis. Nat Methods. 2012;9: 671–675.

69. Guillard RRL, Lorenzen CJ. YELLOW-GREEN ALGAE WITH CHLOROPHYLLIDE C1,2. J Phycol. 1972;8: 10–14.

70. Cornwallis CK, Svensson-Coelho M, Lindh M, Li Q, Stábile F, Hansson L-A, et al. Single-cell adaptations shape evolutionary transitions to multicellularity in green algae. Nature Ecology & Evolution. 2023;7: 889–902.

71. Guillou L, Bachar D, Audic S, Bass D, Berney C, Bittner L, et al. The Protist Ribosomal Reference database (PR2): a catalog of unicellular eukaryote small sub-unit rRNA sequences with curated taxonomy. Nucleic Acids Res. 2013;41: D597–604.

72. Fu L, Niu B, Zhu Z, Wu S, Li W. CD-HIT: accelerated for clustering the next-generation sequencing data. Bioinformatics. 2012;28: 3150.

73. Larsen JB, Larsen A, Bratbak G, Sandaa R-A. Phylogenetic Analysis of Members of the Phycodnaviridae Virus Family, Using Amplified Fragments of the Major Capsid Protein Gene. Appl Environ Microbiol. 2008;74: 3048.

74. Hoops HJ, Witman GB. Outer doublet heterogeneity reveals structural polarity related to beat direction in Chlamydomonas flagella. J Cell Biol. 1983;97: 902–908.

75. Richardson KC, Jarett L, Finke EH. Embedding in epoxy resins for ultrathin sectioning in electron microscopy. Stain Technol. 1960;35: 313–323.

