## Supplemental Figures S1-S13 and Tables S1-S3 for "Latent infection of an active giant endogenous virus in a unicellular green alga"

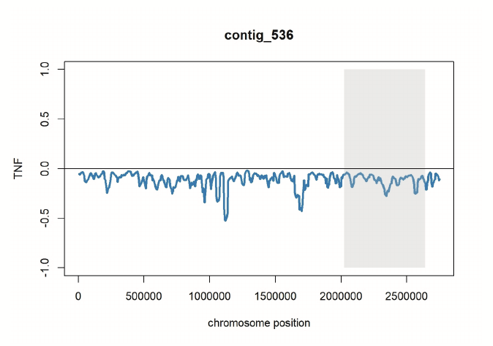

Fig. S1.

**Genome-averaged Tetranucleotide Frequency (TNF) of the viral contig_536.** The GEVE region is shaded in gray.

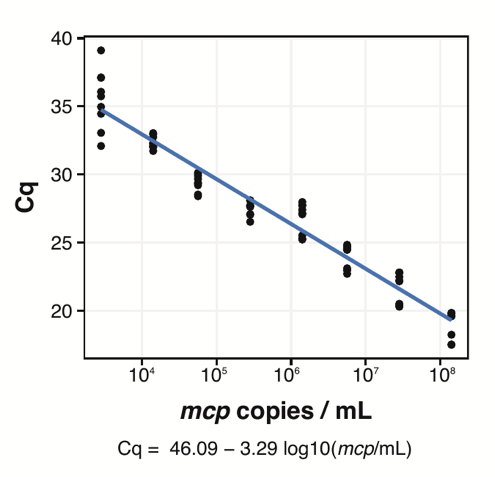

Fig. S2.

**Major capsid protein (MCP) qPCR standard curve.** The curve was calculated from a dilution series ranging from 2.82 x 10^3^ to 1.41 x 10^8^ molecules mL⁻¹ of a synthetic linear oligo containing a fragment of the GEVE *mcp* gene. The plot shows the pooled data from all qPCR runs performed (n=3). The equation for the curve is displayed.

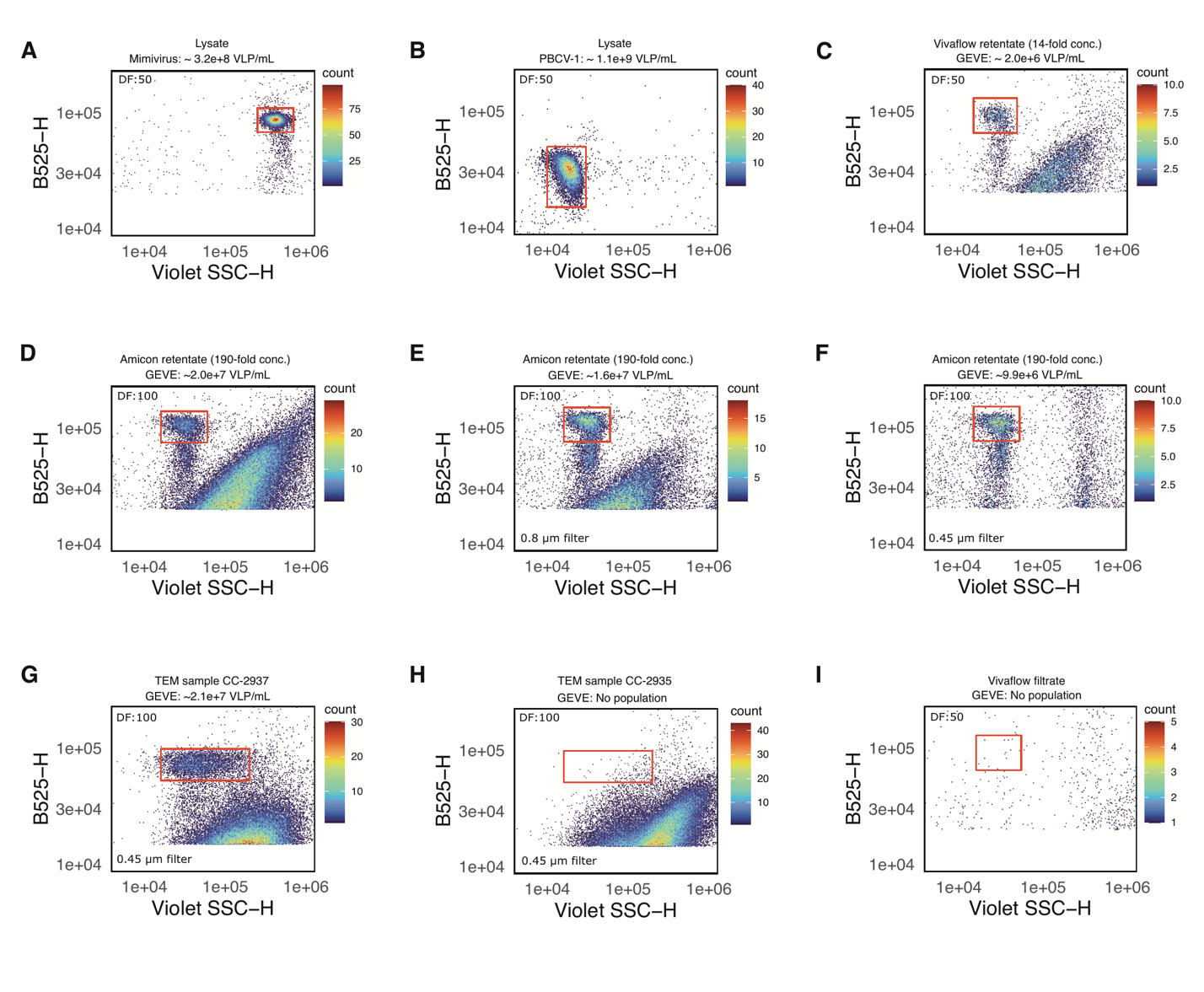

Fig. S3.

**Flow cytograms showing Violet Side Scatter (V-SSC) versus Green Fluorescence (B525) for various large DNA virus lysates and concentrated *Chlamydomonas* *reinhardtii* CC-2937 supernatants.** Samples were fixed with glutaraldehyde and stained with nucleic acid-specific SYBR Safe. The viral abundance (VLPs mL⁻¹) and the dilution factor (DF) before staining are indicated within each plot. **(A)** Mimivirus lysate (positive control). **(B)** Paramecium bursaria chlorella virus 1 lysate (positive control). **(C)** 15-day old CC-2937 supernatant, concentrated 14-fold using a 100 kDa MWCO Vivaflow Tangential Flow Filtration (TFF) unit. **(D)** 15-day old CC-2937 supernatant, further concentrated with an Amicon 100 kDa Ultra Centrifugal Filter (total 190-fold). The measured viral abundance (2.0 x 10^7^ VLPs mL^-1^) was consistent with the expected theoretical abundance after concentrating cultures with 1.6 x 10^5^ VLP mL^-1^ (average calculated by qPCR) 190-fold. **(E)** 190-fold concentrated CC-2937 supernatant filtered through a PES 0.8 µm filter. **(F)** 190-fold concentrated CC-2937 supernatant filtered through a PES 0.45 µm filter. The GEVE population remained detectable, with significant reduction in background noise, likely due to the removal of contaminants and cellular debris. **(G)** 9-day old CC-2937 supernatant, concentrated 40-fold using TFF. A 5.5 mL aliquot was further centrifuged, and the pellet was resuspended in molecular-grade water and filtered through a 0.45 µm PES filter. Sample was used for VLP imaging using negative stain Transmission Electron Microscopy (TEM). **(H)** 9-day old *C. reinhardtii* CC-2935 supernatant, concentrated and visualized as described previously, used for discrimination of false positive VLPs via negative stain TEM. **(I)** Filtrate/permeate from the Vivaflow TFF unit after concentrating CC-2937 supernatants, used as a negative control (contains particles <100 kDa MWCO).

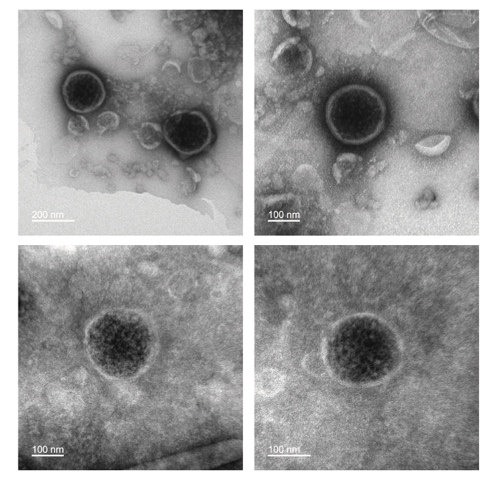

Fig. S4.

**Representative electron micrographs of negatively stained virus-like particles (VLPs).** Concentrated supernatants from *Chlamydomonas reinhardtii* CC-2937 late exponential cultures were prepared using tangential flow filtration and stained with uranyl acetate. The VLPs ranged in diameter from 175 nm to 229 nm, with an average size of 200 nm. Scale bars are provided within each image.

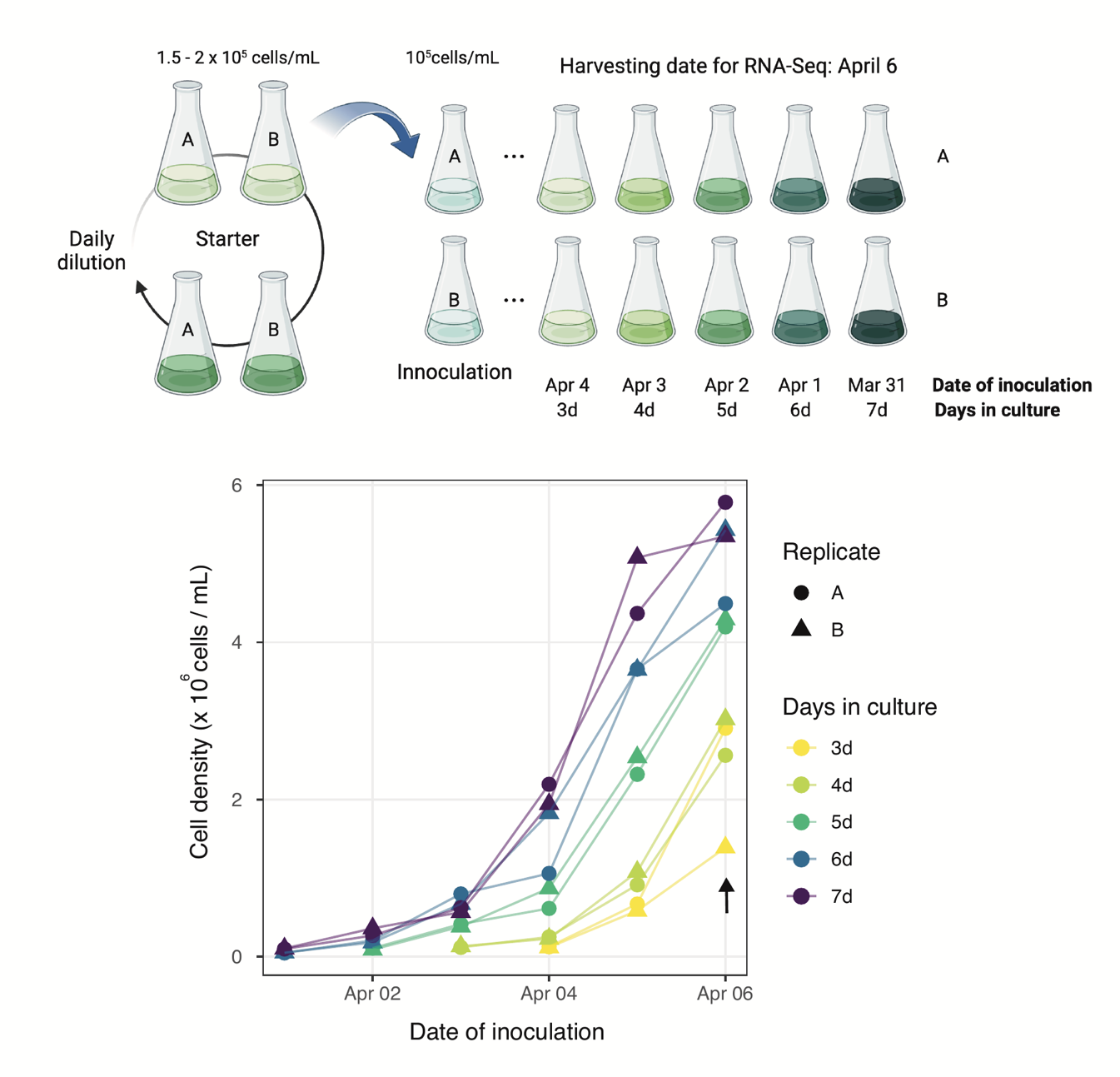

Fig. S5.

**RNA-Seq experiment schematic.** A starter *C. reinhardtii* CC-2937 culture in duplicate was maintained in exponential growth (1.5 - 2 x10^5^ cells mL^-1^) by daily dilution on fresh media. From this starter culture, new flasks were inoculated daily (starting March 31st, 2023), and allowed to grow up to seven days since the start date of the experiment. Following this scheme, all samples were harvested simultaneously on April 6th, 2023 (see arrow in cell density plot), with cultures ranging from 3 to 7 days old, representative of the early to late exponential stages of growth. Measured cell counts are displayed. Experimental design scheme was created with BioRender.com.

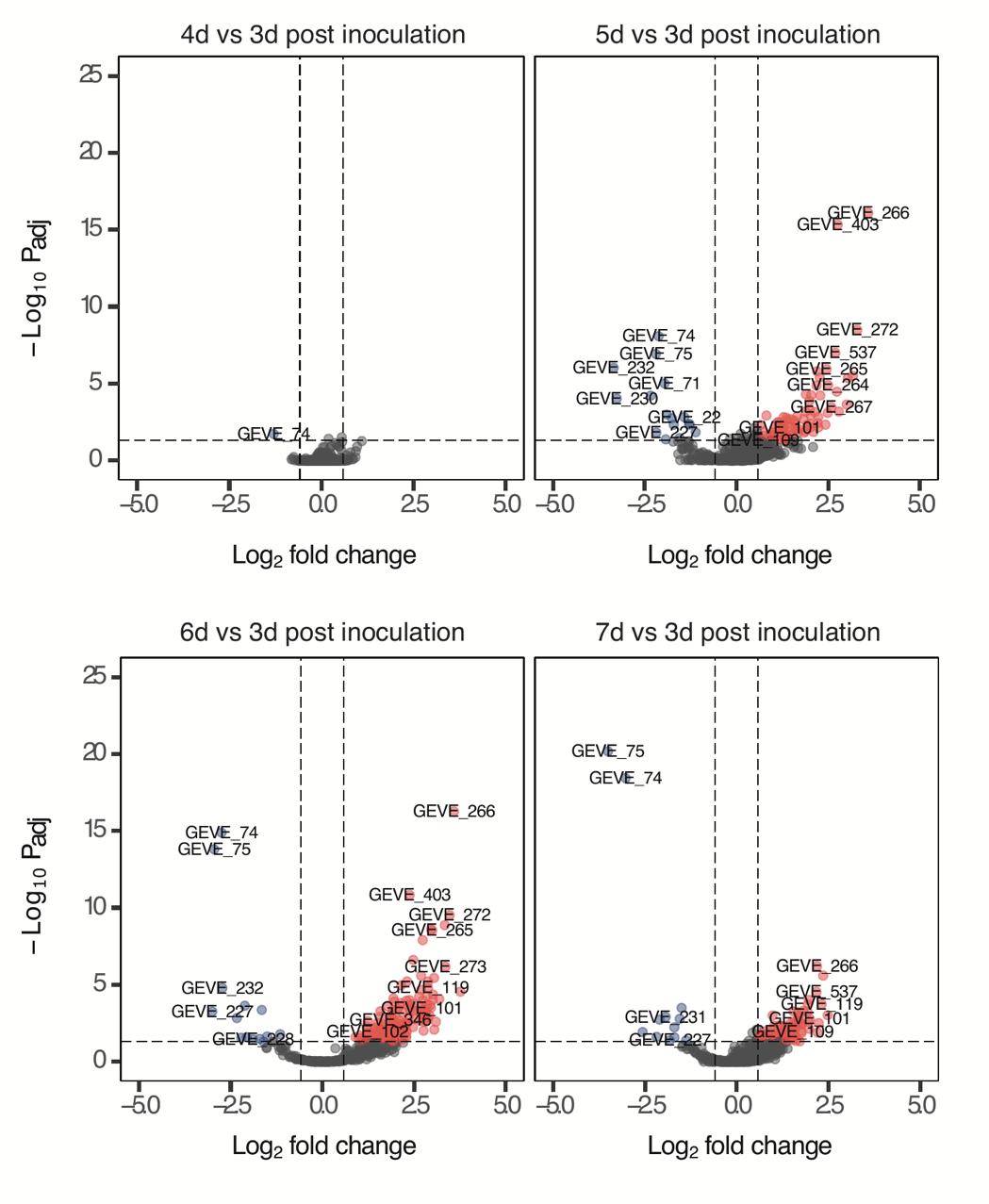

Fig. S6.

**Volcano plots showing GEVE Differentially Expressed Genes (DEGs) at various time points during the growth curve.** Day three post-inoculation (the youngest culture) is used as the reference level for all comparisons. Upregulated DEGs are depicted in red, downregulated DEGs in blue, and non-DEGs (with an absolute Log2 fold change < 1.5 and an adjusted p-value > 0.05) are shown in gray.

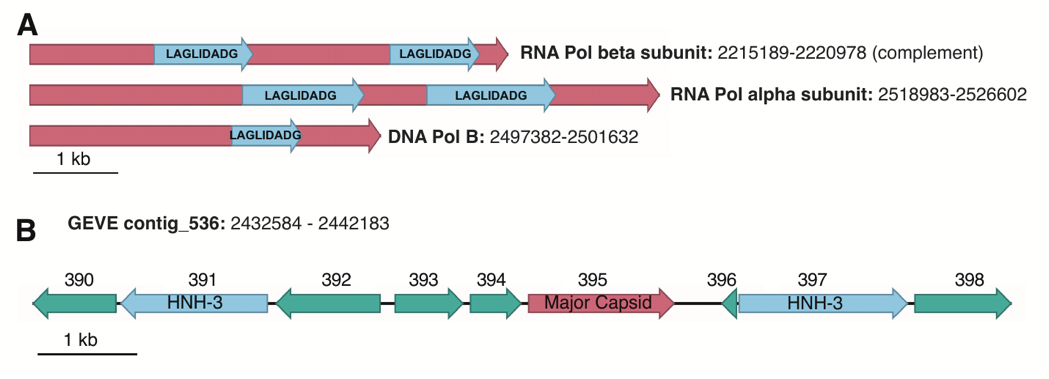

Fig. S7.

**Putative homing endonucleases in the CC-2937 genome.** **(A)** Inteinic LAGLIDADG genes in the GEVE. **(B)** Freestanding HNH-3 nuclease genes in the GEVE. Genes of interest are labeled and the locus nucleotide position in the GEVE contig_536 is shown. The GEVE genes numbers are displayed in (B).

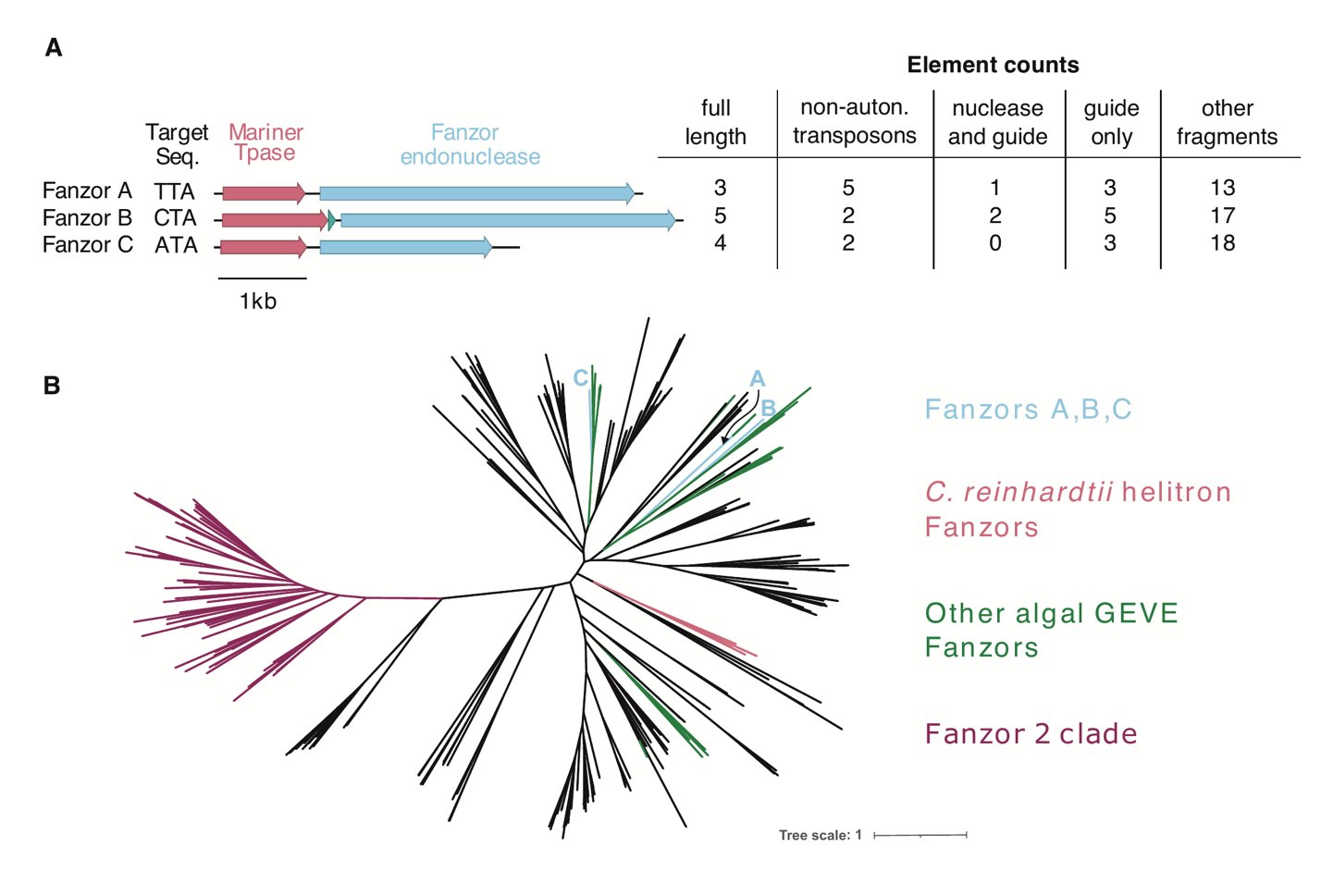

Fig. S8.

**The CC-2937 GEVE encodes three families of Fanzor elements.** **(A)** Diagram of the three families of Fanzor elements found in the CC-2937 GEVE. Consensus complete elements are shown with their predicted gene products labeled. For each family the target sequence and number of elements in various configurations is displayed. **(B)** A phylogenetic tree of Fanzor nucleases. The positions of CC-2937 GEVE Fanzor nuclease and other Fanzor nucleases of particular interest are indicated.

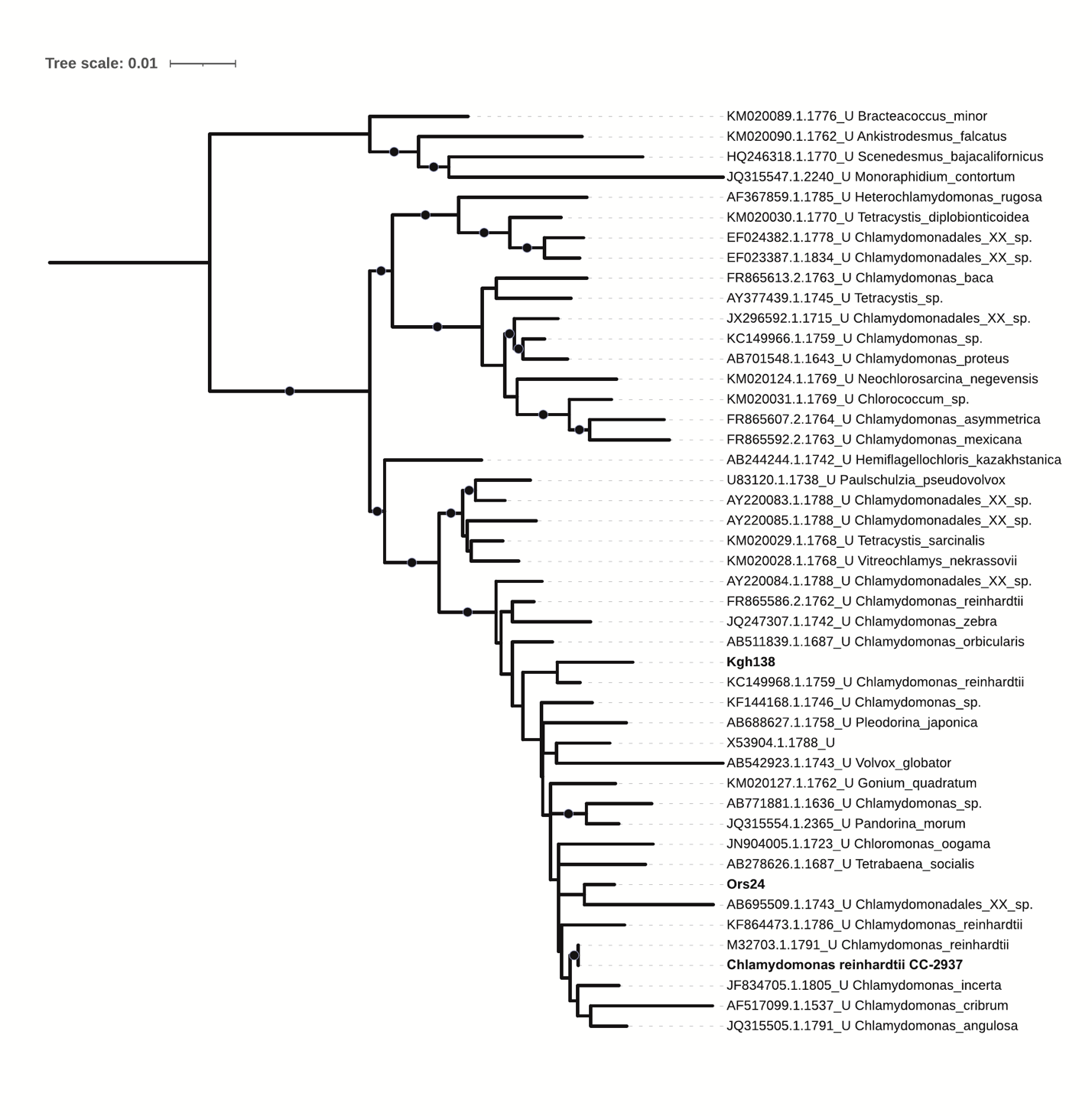

Fig. S9.

**Maximum likelihood phylogenetic tree based on 18S rRNA sequences.** The tree includes sequences from *Chlamydomonas reinhardtii* strain CC-2937 (highlighted in bold), two representative freshwater isolates from Örsjön and Krageholmssjön (Ors24 and Kgh138, also highlighted in bold), and selected sequences from the PR2 database.

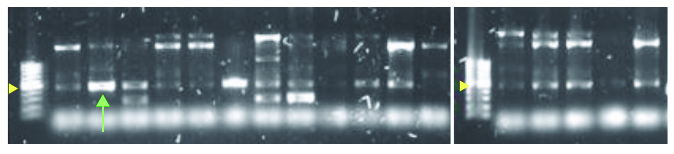

Fig. S10.

**Representative gel of the PCR screen of viral MCPs in *Chlamydomonas* sp. freshwater isolates.** Degenerate primers targeting large algal virus *mcp* sequences (*73*) were used for screening. The yellow arrowhead in the molecular ladder indicates 500 bp. The green arrow points to the band of interest.

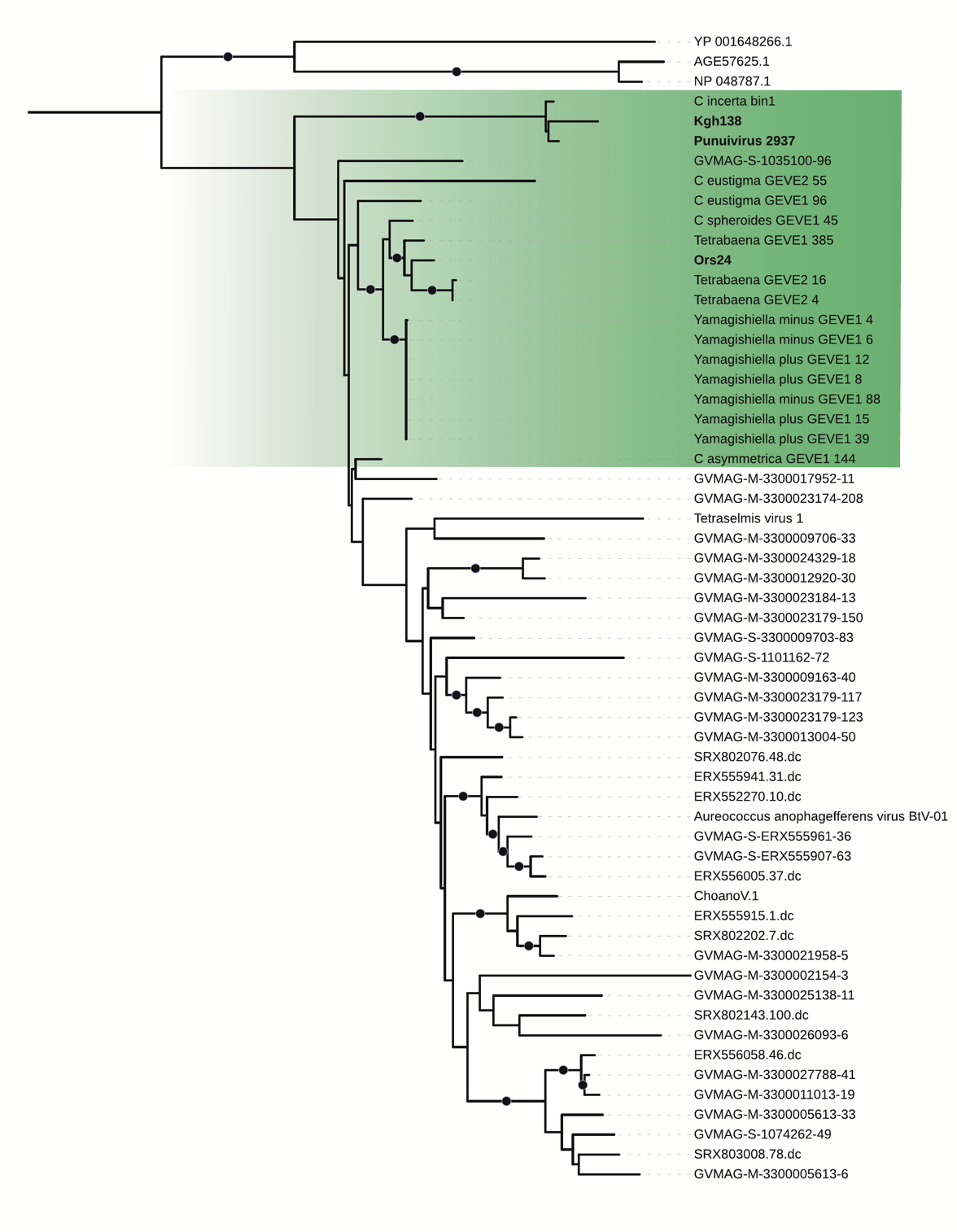

Fig. S11.

**Maximum likelihood phylogenetic tree of viral Major Capsid Protein (MCP) sequences.** The tree includes sequences from *Chlamydomonas reinhardtii* strain CC-2937 GEVE (Punuivirus 2937; highlighted in bold), two representative *Chlamydomonas* sp. freshwater isolates from Örsjön and Krageholmssjön (Ors24 and Kgh138, also highlighted in bold), as well as other representative GEVEs and members from various NCLDV families. The green-shaded area highlights a clade of GEVEs identified in green algae. The name "Punuivirus" is derived from the Incan deity Puñuy, associated with dreams and sleep in Andean cosmovision.

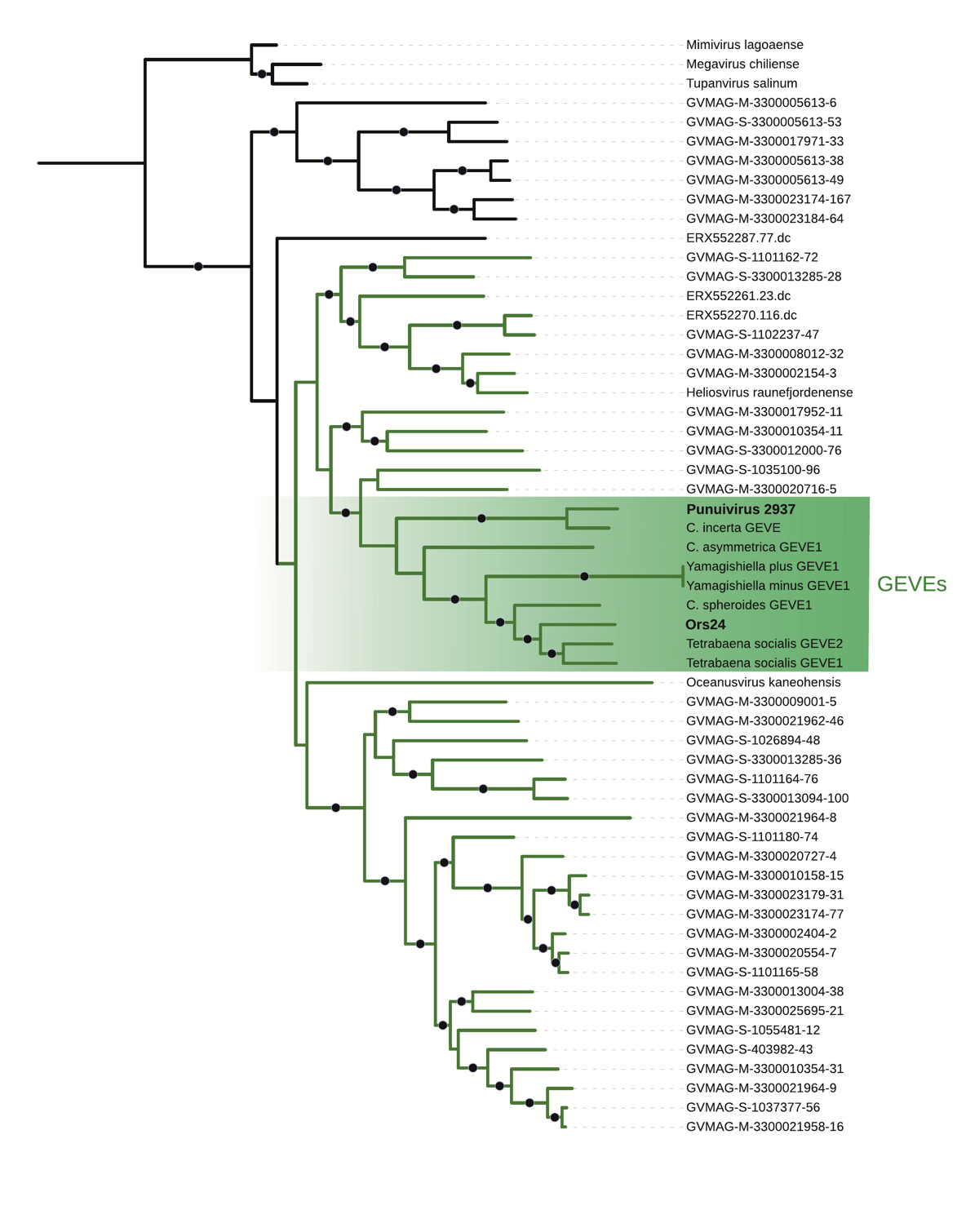

Fig. S12.

**Maximum likelihood phylogenetic tree of viral DNA Polymerase B (PolB) sequences.** The tree includes sequences from *Chlamydomonas reinhardtii* strain CC-2937 GEVE (Punuivirus 2937; highlighted in bold), a representative *Chlamydomonas* sp. freshwater isolate from Örsjön (Ors24, also highlighted in bold), along with other representative GEVEs and members from various NCLDV families. The green-shaded area indicates a clade composed of various GEVEs. The name "Punuivirus" is inspired by the Incan deity Puñuy, associated with dreams and sleep in Andean cosmovision.

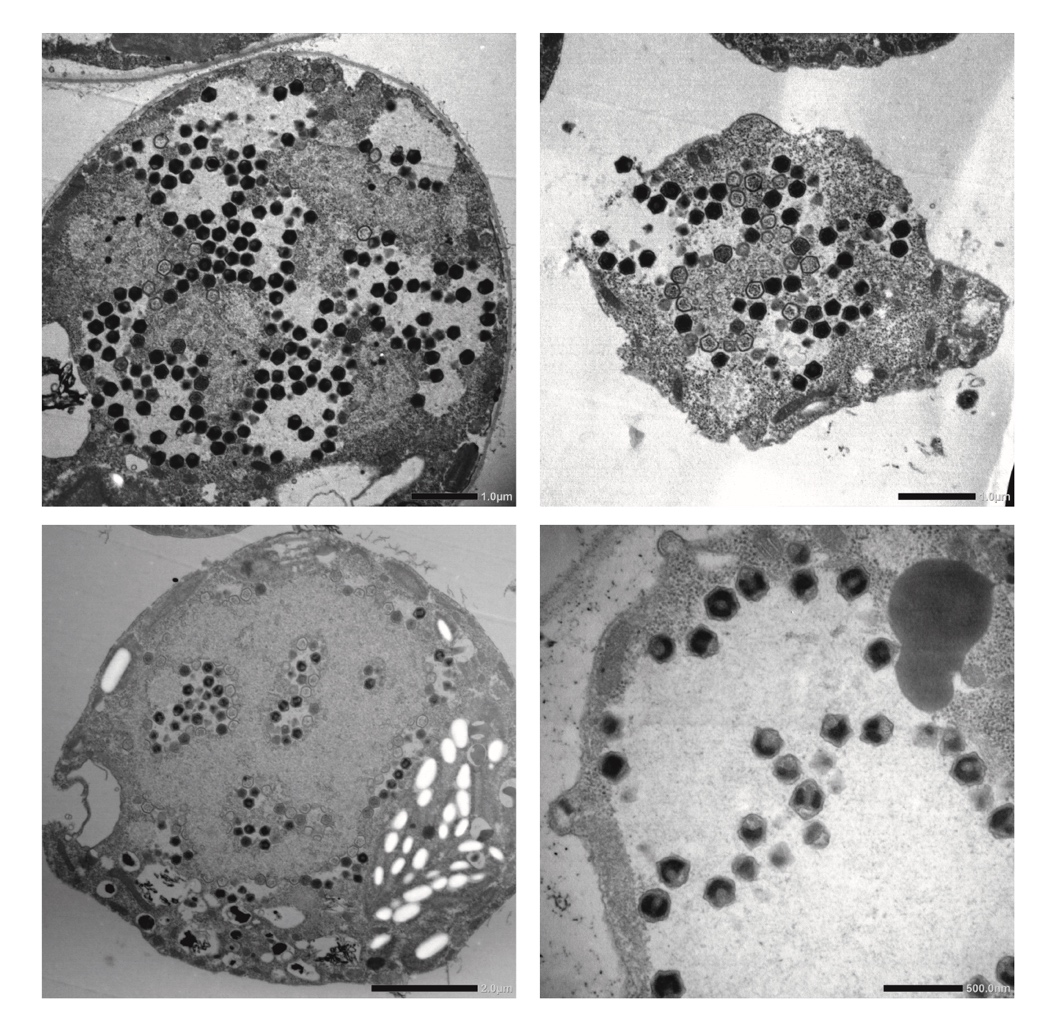

Fig. S13.

**Transmission electron micrographs of ultra-thin sectioned Ors24 cells.** The images show icosahedral virus-like particle (VLP) inclusions formed within the cells. Scale bars are provided within each image.

Table S1.

**Distribution of Terminal Inverted Repeats (TIRs) across *C. reinhardtii* genomes.** TIRs instances of endogenized Punuivirus cr2937 were screened across the strains CC-2937, CC-4532, CC-1952, and CC-2931. Where the TIR sequence of a specific endogenization event is shared by more than one genome, only coordinates for the reference genome (CC-4532) are provided.

| **strain** | **chromosome** | **start** | **end** | **length** | **TSD** | **Notes** |
| --- | --- | --- | --- | --- | --- | --- |
| CC-2937 | contig_536 | 2023525 | 2640528 | 617003 | GCCACG | equivalent to chromosome_15, see main text; full-length Punuivirus cr2937 |
| CC-2937 | contig_437 | 3767614 | 3815820 | 48206 | TCGATC | equivalent to chromosome_16, see main text |
| CC-2937 | contig_337 | 1722678 | 1727950 | 5272 | TACGCC | equivalent to chromosome_07, see main text |
| CC-4532 | chromosome_15 | 987207 | 988297 | 1090 | TGCTTG | termini of left and right TIRs present with TSD |
| CC-4532 | chromosome_09 | 3398202 | 3399030 | 828 | CGGAGC | deletion has almost entirely removed left TIR, TSD present |
| CC-4532 | chromosome_14 | 166841 | 167062 | 221 | NA | one TIR only; present among all strains |
| CC-4532 | chromosome_10 | 3582465 | 3582601 | 136 | NA | one TIR only; centromeric |
| CC-4532 | chromosome_16 | 3581731 | 3581974 | 243 | NA | one TIR only; centromeric; present among all strains |
| CC-1952 | chromosome_15 | 3383809 | 3394843 | 11034 | GCCGTC | termini of left and right TIRs present with TSD |
| CC-1952 | chromosome_13 | 4586079 | 4587151 | 1072 | NA | one TIR only |

Table S2.

**PCR Screening of the *mcp* gene in freshwater *Chlamydomonas* sp. Isolates.** Isolates from two lakes in Southern Sweden, Örsjön and Krageholmssjön, were screened using degenerate primers designed to target MCP sequences of large algal viruses (*73*). Asterisks denote representative isolates from each lake. For the Krageholmssjön isolates, only one isolate (Kgh138) yielded a purifiable and sequencable mcp product. In contrast, multiple isolates from Örsjön were successfully sequenced, and the consensus sequence was identical to that of the representative isolate, Ors24.

| **Number** | **Location** | **ID** | ***mcp* amplification** | **Sequenced** |
| --- | --- | --- | --- | --- |
| 1 | Krageholmssjön | Kgh130 | - |  |
| 2 | Krageholmssjön | Kgh131 | - |  |
| 3 | Krageholmssjön | Kgh132 | positive (faint) |  |
| 4 | Krageholmssjön | Kgh134 | positive (faint) |  |
| 5 | Krageholmssjön | Kgh135 | positive (faint) |  |
| 6 | Krageholmssjön | Kgh136 | positive (faint) |  |
| 7 | Krageholmssjön | Kgh137 | positive (faint) |  |
| 8 | Krageholmssjön | Kgh138 | positive | Yes* |
| 9 | Krageholmssjön | Kgh139 | - |  |
| 10 | Krageholmssjön | Kgh140 | positive |  |
| 11 | Krageholmssjön | Kgh141 | positive (faint) |  |
| 12 | Krageholmssjön | Kgh142 | positive (faint) |  |
| 13 | Krageholmssjön | Kgh144 | positive (faint) |  |
| 14 | Krageholmssjön | Kgh146 | - |  |
| 15 | Krageholmssjön | Kgh147 | - |  |
| 16 | Krageholmssjön | Kgh149 | - |  |
| 17 | Krageholmssjön | Kgh150 | - |  |
| 18 | Krageholmssjön | Kgh151 | - |  |
| 19 | Krageholmssjön | Kgh152 | positive (faint) |  |
| 20 | Krageholmssjön | Kgh155 | positive (faint) |  |
| 1 | Örsjön | Ors09 | positive (faint) | Yes |
| 2 | Örsjön | Ors10 | positive | Yes |
| 3 | Örsjön | Ors11 | positive | Yes |
| 4 | Örsjön | Ors12 | positive | Yes |
| 5 | Örsjön | Ors13 | - |  |
| 6 | Örsjön | Ors14 | - |  |
| 7 | Örsjön | Ors15 | positive | Yes |
| 8 | Örsjön | Ors16 | positive | Yes |
| 9 | Örsjön | Ors17 | positive |  |
| 10 | Örsjön | Ors18 | - |  |
| 11 | Örsjön | Ors20 | positive | Yes |
| 12 | Örsjön | Ors21 | positive |  |
| 13 | Örsjön | Ors22 | - |  |
| 14 | Örsjön | Ors23 | positive (faint) |  |
| 15 | Örsjön | Ors24 | positive | Yes* |
| 16 | Örsjön | Ors25 | positive |  |
| 17 | Örsjön | Ors26 | - |  |
| 18 | Örsjön | Ors27 | positive | Yes |

Table S3.

RiboZero Plus Supplemental Probes used for rRNA depletion.

| **Name** | **Sequence (5' - 3')** |
| --- | --- |
| pool_probe_1 | AGGGACGTAATCAACGCGAGCTGATGACTCGCGCTTACTAGGCATTCCTC |
| pool_probe_2 | GACAGTGAAGCCCAGGAGCCCGTCCCCGGCAGGAAGGTGGAGCAGAGCAG |
| pool_probe_3 | GACCGCACCACCCCACCCGAAATCCAACTACGAGCTTTTTAACTGCAACA |
| pool_probe_4 | CTCTCAATCTGTCAATCCTTCCCGTGTCTGGACCTGGTAAGTTTCCCCGT |
| pool_probe_5 | ACTCCCCCCGGAACCCAAAAACTTTGATTTCTCATAAGGTGCTGGCAGAG |
| pool_probe_6 | TATCCGTTGTTGAGAGTTGTCTTGGTTAGTGAGCACTAGCTCTAAGCTAG |
| pool_probe_7 | GACAAGCACAGCTCCAAGCCATCTACTAAAGCATTAGCAACGAGTTGCCT |
| pool_probe_8 | ACCGAGACAGCCAGGTCCGCTCTTGCTCCAAACATGTTGGAGTAGGGCGA |
| pool_probe_9 | TTGCAATAATCTATCCCCATCACGACGCGGTTTACAAGATTACCCGGGCC |
| pool_probe_10 | TCGTTAAGGGATTTAGATTGTACTCATTCCAATTACCAGACGCGAAGCGC |
| pool_probe_11 | GTAAATCCAAGAATTTCACCTCTGACAACGGAATACGAATGCCCCCGACT |
| pool_probe_12 | TCTCCTTCCTCTAGGTGGGAGGGTTTAATGAACTTCTCAAGCAAGGCCAG |
| pool_probe_13 | GGTAGCCATCTCTCAGGCTCCCTCTCCGGAATCGAACCCTAATCCTCCGT |
| pool_probe_14 | GGGGAATCCTTGTTAGTTTCTTTTCCTCCGCTTATTGATATGCTTAAGTT |
| pool_probe_15 | TACTCCGGTCCCACAGACCAACAGATAGGCCAGAGTCCTATCGTGTTATT |
| pool_probe_16 | TTAGTTTTGGTTTGGTGTTGGTTGGTTAGTATAAGAGACCGATGCCTTGC |
| pool_probe_17 | CCTGAGCTCAGGTCGAGAAAATAGGGGTTTGCTTCGGCACAGGGCCAAAG |
| pool_probe_18 | ATAAGCGGGTCCATGCATCAACCTCTGTACTTCAGCTGACCCGGCGTCTG |
| pool_probe_19 | TGCGATGCTGACTATTTAGCAGGCTGAGGTCTCGTTCGTTACCGGAATCA |
| pool_probe_20 | CACAGTATAAGCAGTTTATACTTAGACATGCATGGCTTAATCTTTGAGAC |
| pool_probe_21 | TTATCGCCTCATACTTCCATTGGCTAAACGCCAATAGTCCCTCTAAGAAG |
| pool_probe_22 | TATCACCCTCTCTGACGCGGCATTCGATCCGACTTGAGTTCCGCCAGCCC |
| pool_probe_23 | GTATTTCTGCAATTCACACTACGTATCGCATTTCGCTGCGTTCTTCATCG |
| pool_probe_24 | TTATCTAATAAATACGACCCTTCCAGAAGTCGGGTTGTGCGCACGTATTA |
| pool_probe_25 | TGAGGCAGACATGCTCTTGGCCGAAGCCTCGAGCGCAATATGCGTTCAAA |
| pool_probe_26 | TGGTAGGCCTCTATCCTACCATCGAAAGTTGATAGGGCAGAAATTTGAAT |
| pool_probe_27 | CGGTTATCCGAGTAGTAGGTACCATCAAATAAACTATAACTGATTTAATG |
| pool_probe_28 | CTCAATCCGAACACTTCACCAGCACACCCAATCGGTAGGAGCGACGGGCG |
| pool_probe_29 | CTGCCAATCCCTAGTCGGCATCGTTTATGGTTGAGACTACGACGGTATCT |
| pool_probe_30 | GCGTAACTTTTACATTGCGCCCAGACCTGGCCCGTTTGCTTTTACACAGA |
| pool_probe_31 | CAGAGCGTAGGCCTGCTTTGAACACTCTAATTTACTCAAAGTAACCTCGC |
| pool_probe_32 | CGAGCCCATACGATTCGTGAAGTTATCATGATTCACCGCAGGTCGGGCAG |
| pool_probe_33 | GGCTCGTTGGTCGCGTCAGTGTAGCGCGCGTGCGGCCCAGAACATCTAAG |
| pool_probe_34 | CCCAACTTTCGTTCTTGATCAATGAAAGTATCCTTGGCAAATGCTTTCGC |
| pool_probe_35 | TAGACTACAATTCTCCAAAGGGAGATTTTCAAGTTGGGCTATTCCCGGTT |
